# Nineteenth-century emergence of a seafloor ecosystem beyond late Quaternary limits

**DOI:** 10.64898/2026.04.26.720923

**Authors:** Sara S. Kahanamoku, Ivo A.P. Duijnstee, Ingrid L. Hendy, Richard D. Norris, Seth Finnegan

## Abstract

Mismatches between ecological time and geologic time complicate our ability to effectively compare contemporary ecosystem changes with those of the past. We amassed sub-decadal paleoecological data from the California Current System (CCS) to examine how benthic ecosystem structure evolved over the past 34,000 years in the Santa Barbara Basin across the glacial-to-interglacial transition. Our results indicate that even deep-sea ecosystems of the anthropocene *sensu lato* – the past few centuries characterized by outsized human impacts – are distinct compared to any point in the last 34,000 years. Novel ecosystem states and heightened variability began in the early 1800s AD, suggesting that colonial-era human land-use drove the emergence of a novel ecosystem state more than a century before global climate warming began in earnest.

## The invisible timescales of global change

Novel human activity has caused the oceans to warm faster than at any point in the last 2,000 years (*1–3*), driving exponential losses in dissolved oxygen (O_2_) concentrations since at least the 1950s (*4*) that have altered the distribution and behavior of marine organisms across the globe (*5*). As ocean warming and deoxygenation is projected to further accelerate into the next century and beyond (*6*), understanding how marine ecosystems have responded to past rapid environmental changes is critical for accurately predicting ecosystem futures.

Yet the timescales of anthropogenic impacts pose a fundamental challenge: they are uniquely rapid but have been occurring for at least several centuries, meaning that most modern ecosystems have likely already undergone significant alteration (*5, 7–10*). As parts of the Earth system respond to environmental change at varying rates and timing due to the combined effects of climate warming and anthropogenic modification, there is increasing concern about whether, when, and how system-scale tipping points might be reached (*11*). Uncertainties about the rate, magnitude, and nonlinear interactions of global changes over the past several decades – popularly referred to as the “Anthropocene” (*12, 13*) – complicate finding definitive answers to important, longstanding questions, such as whether modern biodiversity losses can truly be considered a “sixth mass extinction” (*14–16*), what represents an ecological tipping point (*17, 18*), and the timescales over which we can expect novel ecosystems (*19*) to emerge under future climate scenarios (*20, 21*). While we have short-term, highly resolved records of ecological change from direct observations (*22*) and over multi-millennial geologic timescales (*23*), we often lack the intermediate-resolution records that link these disparate scales (*10, 24*). Thus, integrating the highly-resolved ecosystem changes of the Anthropocene with the long-term behavior of ecosystems captured in the fossil record requires exceptional systems that can span across the invisible timescales of global change.

### Placing the anthropocene in geologic context

Here, we use a 34,000-year-long composite sediment core record from the Santa Barbara Basin (SBB) of Southern California (**SI Figure 1**) to place contemporary ecosystem states in geologic context. The SBB is an ideal natural laboratory in which to link the patterns and scales of ecological responses to human and climate impacts through time. First, while the proximity of the SBB to the California coast makes it prone to indirect impacts of human activity, benthic ecosystems in the basin’s center lie in deep-ocean water depths (>580m) that provide a buffer for direct effects, limiting the types of stressors to the system and facilitating deconvolution of compounding effects. Second, high productivity in the overlying California Current System (CCS) creates persistent hypoxia that preserves millimeter-scale seasonal varve couplets that record climate, hydrographic, and biological variability within the broader CCS over the past tens of thousands of years at an unusually fine temporal resolution (*25–30*). Third, the sub-decadal resolution of this paleo archive can be readily combined with information from several long-term monitoring programs, whose continuous operation since the 1950s AD makes the CCS one of the best-observed marine ecosystem complexes in the world (*31, 32*). As a result, the SBB is one of the few marine systems where direct modern observations can be coupled with high-resolution paleontological and paleoclimate data to retroactively extend ecosystem monitoring, evaluate the rate and drivers of past ecological change, and determine how anthropogenic change may be driving the emergence of novel ecosystems.

Hypoxic conditions have persisted in the central SBB since at least the late Pleistocene (*28, 29, 33*), leading to the development of benthic environments where macrofauna are largely excluded (*34, 35*) Benthic foraminifera comprise the majority of eukaryotic biomass in the deep SBB, both in the present day (*36, 37*) and over geologic time (*34, 38, 39*). As a result, benthic foraminifer compositional data broadly reflect the status of the SBB benthic ecosystem over millennia (*34, 38*) (**SI Text**). Our time series of benthic ecosystem structure in the SBB reaches back into the late Pleistocene, capturing ecological responses to rapid environmental fluctuations across several glacial interstadial events (GI6 to GI2.1: ∼33 ka, 32 ka, 30 ka, 28 ka, 27 ka, and 23 ka) and the glacial termination (GI-1: ∼18-11 ka) (*40*). During these periods of abrupt change, sea surface warming of 4 to 7°C (*41, 42*) is considered to have occurred over decades to centuries (*43–45*), driving concurrent rapid changes to ecological community structure at all ocean depths (*30, 34, 38*). As Pleistocene abrupt warming is similar in magnitude (but not absolute value) to late 21^st^ century climate projections (*46*) and may be similar in rate to present-day warming (*45*), these events may hold critical information for presaging future climate impacts.

We combined previously published benthic foraminifer count records (core MD02-2503, ∼7-yr sample resolution) (*38, 39, 47*) with a new record (MV1012-BC1, ∼2.2 yr sample resolution) (*48*) to create a composite dataset of community composition from 34 kya to 2008 AD (**SI Table 1**). We estimated benthic foraminifer accumulation rates by normalizing counts to the time and sediment volume in each sample following standard practice. To represent environmental change through time, we compiled paleoceanographic proxies for sea surface temperature (alkenone saturation and planktic foraminiferal δ^18^O) (*41, 42, 49, 50*), source-water oxygenation (δ^18^N) (*51, 52*), sedimentary redox conditions (trace-metals Re/Mo, Mo/Al) (*53–55*), organic matter export (TOC) (*51, 53, 56*), sediment flux (mass accumulation rate) (*57, 58*) from twelve sediment cores sampled at or near the same locality within the SBB (**SI Figure 1; SI Table 1**) and placed all data within a unified temporal framework (see **SI Text**).

### Emergence of an anthropocene ecosystem distinct from glacial-interglacial states

A non-metric multidimensional scaling ordination (NMDS with euclidean distance on log-transformed benthic foraminifer accumulation rates; see **SI Text**) indicates that SBB benthic ecosystems experienced repeated faunal turnover correlated with cyclic oscillations in global temperature and marine deoxygenation over the past 34,000 years (**Figure 1A**). The largest turnover events (which in our dataset can be identified by the greatest distances in the NMDS in **Figure 1A**) correspond with episodes of rapid or extreme warming (**Figure 1B; SI Video 1**). Community changes are largely synchronous with globally-recognized Pleistocene glacial interstadial events and Holocene climate anomalies (hereafter, “abrupt events”) (*40*) and their regional expressions within the California Current System (*30, 59*). Communities from abrupt events occupy a shared region within ordination space (hereafter, “ecosystem space”) (*60*), suggesting that rapid warming produced consistent responses in benthic foraminifer community structure despite wide variation in background climate conditions in the CCS from the last glacial period to the present.

**Figure 1:**
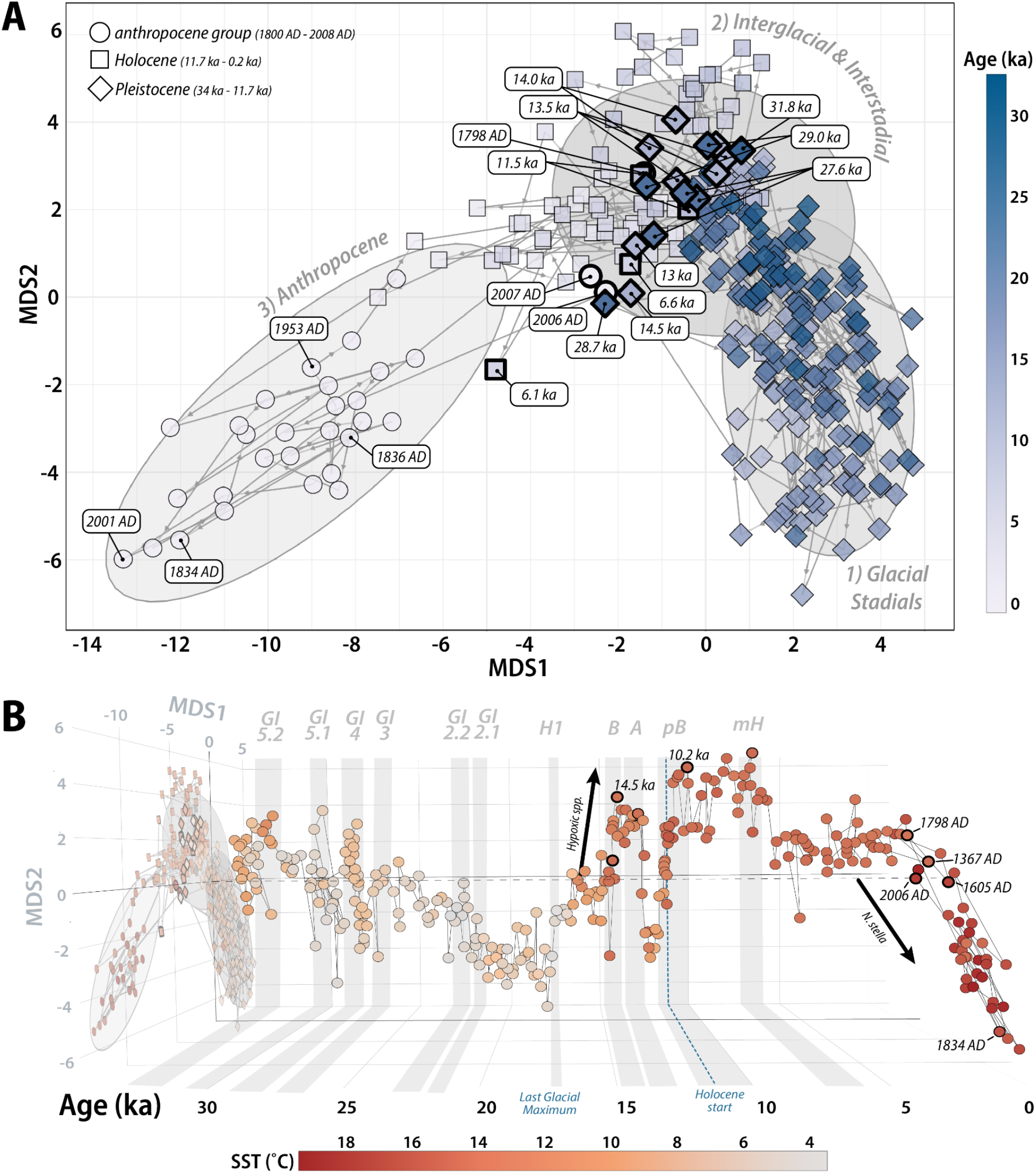
34,000 years of SBB benthic community composition. (**A**) NMDS ordination with k-means cluster groupings (best-fit k = 3) of Euclidean distances between foraminiferal assemblages (time- and volume-normalized, log-transformed counts: see **SI text**). While assemblages largely group by age, several samples from the 19^th^ and 20^th^-century group most closely with abrupt interstadial and deglacial warming events. Points denote foraminiferal assemblages from different time intervals, obtained from two cores, MD02-2503 and MV1012-BC1. Samples from the Pleistocene (in our data, 34 ka - 11.7 ka) are represented by rhomboids; Holocene (here, 11.7 ka to ∼1800 AD) by squares, and “anthropocene” (*sensu lato*; here, 1800 AD to 2008 AD) by circles. (**B**) NMDS ordination for benthic foraminiferal assemblages, with MDS axes 1 and 2 shown plotted against time, 34 ka to present. Grey bars indicate interstadial events expressed in the SBB (*30, 59*) that correspond with Greenland Interstadial events from the INTIMATE ice core stratigraphy (*40*): GI-5.2 and GI-5.1, with peaks at 32.5 and 30.8 ka; GI-4 at 28.9 ka; GI-3 at 27.7 ka; GI-2.2 and GI-2.1 at 23.3 and 23.0 ka; Henrich Event 1 (H1) at 17.5 ka (*84*); the initial Bølling warming (B), GI-1e, at 14.6 ka; Allerød warming (A), GI-1a at 13 ka; pre-Boreal warming (pB) from 11.5-10 ka (*41, 85*); and the mid-Holocene (mH) warming from 6.5-6 ka (*86*). Assemblages with the largest magnitude of change in a 2D ordination space, which broadly correspond with abrupt warming events, are labeled with their respective ages. Fill color in (**A**) represents sample age in thousands of years (ka), while (**B**) represents sea surface (SST) estimates from alkenone (*49, 50*) and oxygen isotope proxy data (*41, 42*) and direct SST measurements from the CalCOFI monitoring program (https://calcofi.org).

A best-fit k-means cluster analysis resolved three community composition clusters: glacial stadial (GS) communities, common during the cooler stadial periods of the last glacial; interglacial and glacial interstadial (IG/GI) communities, common during warm glacial events (interstadials) and following the last glacial termination; and a third group we consider to represent anthropocene communities, common following 1798 AD (**SI Table 2**). We characterized each cluster group based on ^14^C chronostratigraphic ages (*30, 39*), updated to reflect regional ΔR (the marine reservoir offset) (*26*) and global climate event stratigraphy (*40*), and included estimates of past sea surface temperature (SST) (*41, 42, 49, 50*) to identify the broad paleoenvironmental conditions represented in each cluster. In spite of relatively high within-group variability, communities typically remain within the ecosystem space occupied by their respective cluster (**Figure 1A**), indicating that GS, IG/GI, and anthropocene communities represent three distinct ecosystem states. As noted previously, abrupt events are an exception to this trend: Pleistocene-age communities from intervals of peak interstadial warming often (but not always) move to a region of ecosystem space occupied primarily by post-glacial Holocene faunas. These similarities suggest that rapid interstadial warming may have presaged the climatological and ecological transition of the early Holocene.

In contrast, there is almost no overlap between anthropocene communities and those from IG/IS and GI groups. A notable exception is the two latest timepoints from our core record, dating to 2007 and 2008 AD, which group closely with communities from several abrupt events (GI-4: 28.7 ka; GI-3: 27.6 ka; Bølling: 14.5; Allerød: 13 ka, and the mid-Holocene Climatic Optimum: 6.6 ka; **Figure 1A**). Ecological similarities between the late 2000s and Pleistocene-to-Holocene abrupt events are likely a reflection of severe anoxia in the deep SBB during these intervals (**SI Figure 2**). For example, rapid warming and associated oceanographic changes across the deglacial transition drove prolonged benthic anoxia (*47, 61, 62*) that resulted in low population densities for benthic foraminifera, and in some cases caused decades-long faunal die-offs in the deep SBB (captured in our data during the Bølling at 14.4 ka and pre-Boreal at 11.6 ka) (*39*). Low population densities in the late 2000s likely reflect rapid deoxygenation of the CCS between the 1990s and 2000s, when SBB bottom-waters experienced O_2_ losses of nearly 20% over a single decade (*63*).

Deglacial and late-2000s benthic communities also share similarities in the rise in abundance of the anoxic extremophile taxon *Nonionella stella. N. stella*’s high metabolic plasticity and ability to respire and generate energy under dark euxinic conditions (*36, 64–66*) allows this taxon to proliferate during severe deoxygenation events (*37*). A more than 100-fold increase in *N. stella* accumulation rates (reflecting *N. stella* population density; see **SI Text** for discussion) from 1950 AD through the late 2000s (**SI Figure 3**) indicates that its current status as the dominant taxon in the SBB (*36, 37*) may be an anthropocene phenomenon. Indeed, NMDS loadings show that anthropocene communities are largely characterized by the increasing influence of *N. stella* (**SI Figure 2**). Prior to the past several decades, *N. stella* only reached relative abundances >30% during the rapid warming of the Bølling (∼14.6-14 ka) and pre-Boreal (∼11.5-11.3 ka) events. In both cases, SST in the CCS is thought to have increased by more than 7°C over several centuries or less (*41, 42*). Our data suggest that prolonged anoxia during the late 2000s caused similar spikes in *N. stella* relative abundance followed by a more than 40-fold decline in population density, reflecting similar trajectories to the extended anoxic events of the deglacial transition (**SI Figure 3**).

Irrespective of shared faunal features between the last few decades and past deglacial warming, anthropocene communities as a whole are fundamentally distinct from their Holocene and Pleistocene counterparts. Ecological theory suggests that such an abrupt change – occurring across just three decades between 1798 and 1834 AD – may represent a catastrophic shift to a contrasting alternative stable state (*67, 68*). To represent the magnitude of change in community composition through time, we used the euclidean distances between samples underpinning the NMDS in Fig. 1 as a measure of ecological distance (*60*). Large ecological distances (i.e., outliers, or values above 1.96σ) are considered to represent saltatory change: abrupt faunal turnover events characteristic of successional recovery following a major disturbance or, in extreme cases, marking wholesale ecological regime shifts (*60, 69*). We find that anthropocene communities have higher ecological distances on average (d = 5.21, σ = 2.43) than glacial stadial and interglacial/glacial interstadial communities (GS d = 4.20, σ = 1.89; IG/IS d = 4.51; σ = 2.05). Pronounced faunal changes were relatively rare over the past 34,000 years, occurring just twelve times in the past 34,000 years (**SI Table 3**). Between the Pleistocene and latest Holocene, pronounced faunal turnover was constrained to periods of interstadial and deglacial rapid warming (**SI Figure 4**). Following the deglacial transition, no large faunal turnover events occurred until ∼1400 AD, when four of the twelve total faunal turnover events of the past 34 millennia occurred in rapid succession. Two major turnover events have occurred in just the past two centuries, beginning in 1798 AD with the highest ecological distance of the past 34 kyr (**SI Figure 4**). These findings are robust to analytical time-averaging (**SI Figure 5**).

### Human impacts and natural climate variability as 19^th^-century drivers of change

Anthropocene assemblages are not only characterized by high magnitudes of change but are also highly temporally variable (**SI Video 1**). To identify the environmental factors shaping ecosystem states through time, we assess relationships between oceanographic proxy variables representative of bottom-water oxygen availability, sedimentary redox conditions, and depositional flux to the deep SBB (**SI Table 1; SI Figure 6**). Three proxies were significant predictors of both community composition (p < 0.001; **SI Figure 7**) and ecological distance (p < 0.01; **SI Figure 8**): total organic carbon (TOC weight %), representing organic carbon delivery to the seafloor (*70*); bulk sedimentary δ^15^N, reflecting oxygenation of the Eastern Tropical North Pacific, a primary source of intermediate water for the California Current System (*71, 72*); and sedimentary Molybdenum enrichment (Mo/Al, normalized to account for variability in terrestrial input), indicating strongly reducing conditions and high concentrations of free H_2_S in sedimentary pore waters (*73*). TOC is the best predictor of community composition in generalized linear models (R^2^ = 0.74), followed by δ^15^N (R^2^ = 0.51) and Mo/Al (R^2^ = 0.34) (**SI Table 4**). To account for potential nonlinearity and assess the relative importance of predictors, we used a gradient-boosting classification machine trained on the primary NMDS axes of variation (*74*). Likewise, TOC was the best predictor in GBMs (gain = 68%), followed by δ^15^N and Mo/Al (gain = 22% and 10%, respectively; **Figure 2**). While all three variables were predictors for ecological distance, their explanatory power was relatively weak in both GLMs (R^2^ < 0.06) and a linear mixed model with cluster group as a random effect (marginal R^2^ = 0.14; **SI Table 5**).

**Figure 2:**
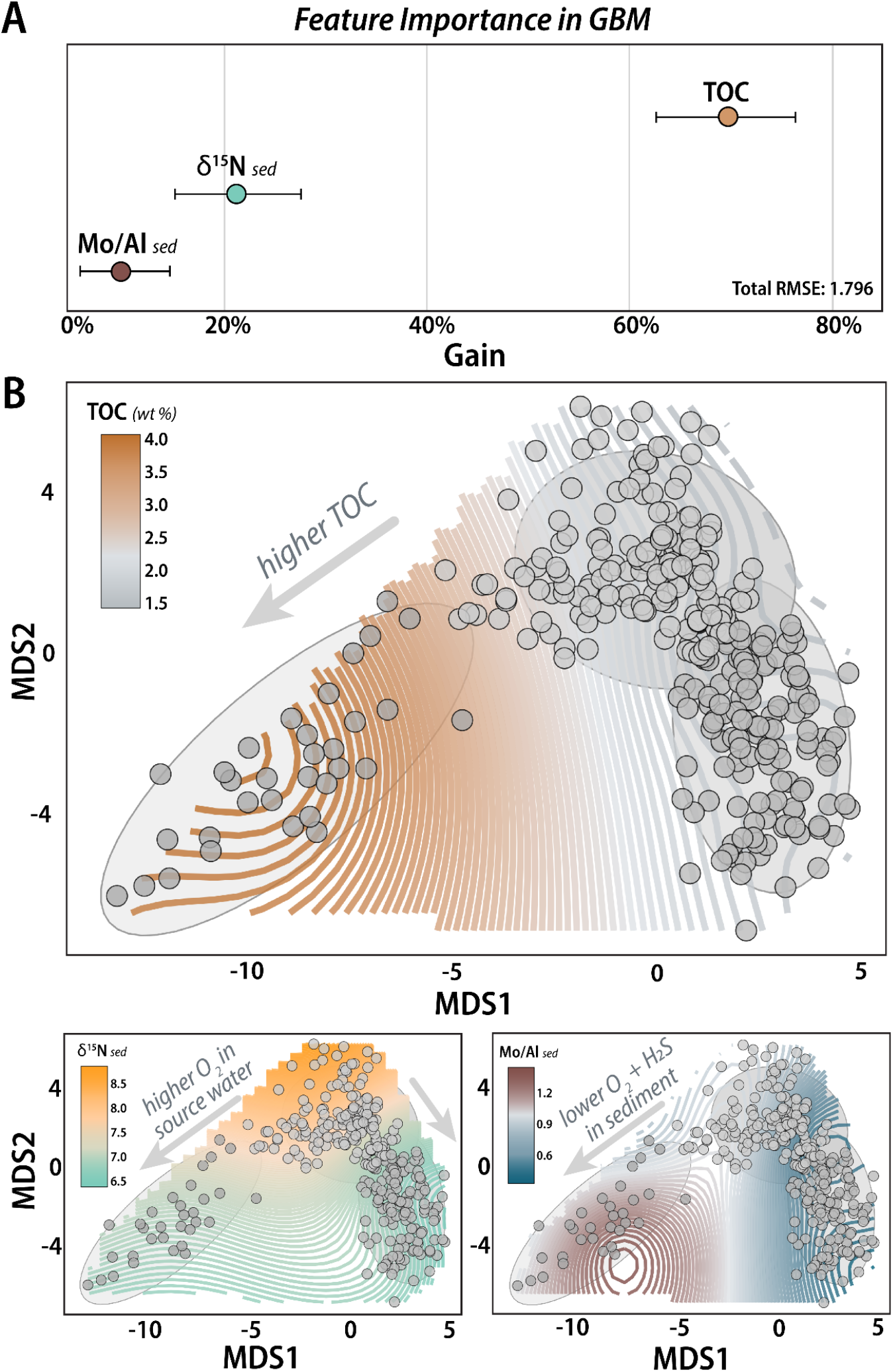
Model performance and gradient visualization for environmental proxy variables significantly predicting community composition. (**A**) Feature importance (expressed as gain) in a Gradient Boosting Machine (GBM) model. Total organic carbon (TOC) explains the highest proportion of variance in community composition (68%), followed by sedimentary δ^15^N (22%), and Mo/Al (10%), Error bars denote 95% bootstrap confidence intervals. (**B**) NMDS ordinations of Euclidean distances between benthic foraminifer community compositions with fitted contours for environmental variables. Panels show, in counter-clockwise order from top, fitted contours for TOC, δ^15^N, and Mo/Al. Points denote community composition at each sampled time point; ellipses denote k-means cluster groups; and coloration of contours denotes value of each fitted variable. Arrows indicate the direction of environmental gradients.

### A new ecological regime in the anthropocene

The timing of ecosystem changes in the early 1800s, nearly a century before climate warming began in earnest (**Figure 3**), points to human land use as a novel factor that may have kickstarted the transition to ecosystem states at a rate exceeding even the glacial-interglacial threshold (*68, 75*). In particular, novel land-use changes during the European colonization of California may be partially responsible for this benthic ecosystem regime shift. Intensive land clearing and livestock grazing across the Mission period (1769-1848 AD) and subsequent American settlement (1848 AD to present) likely drove 5-fold stepwise increases in sediment deposition in the Southern California bight beginning in the mid 1700s AD (*76–78*) (**Figure 3**; **SI Figure 9**). Colonial-era acceleration of sedimentation on the California margin is implicated in a local ecosystem collapse and extirpation event on the Santa Monica shelf in the early 1800s AD (*79*). Similarly, high sedimentation rates may have initiated a life history regime shift in foraminiferal species of the genus *Bolivina* (**Figure 3E**) as well as order-of-magnitude declines in population density among SBB benthic foraminifera (*80*) in the early-to-mid 1800s AD (**Figure 3F**). These concurrent ecosystem changes suggest that the downstream impacts of human land-use change were likely widespread on the California margin in the 19^th^ century. Additional human modification in the 20^th^ century has since caused declining sediment loads within regional catchments and changes to the composition of detrital sediment input to the SBB (*76*). 20^th^ century catchment modifications may explain why sediment input has remained relatively consistent to Southern California’s oceans since the 19^th^ century, despite the intense urban development of greater Los Angeles to its present-day area of more than 85,000 km^2^ (*78*).

**Figure 3:**
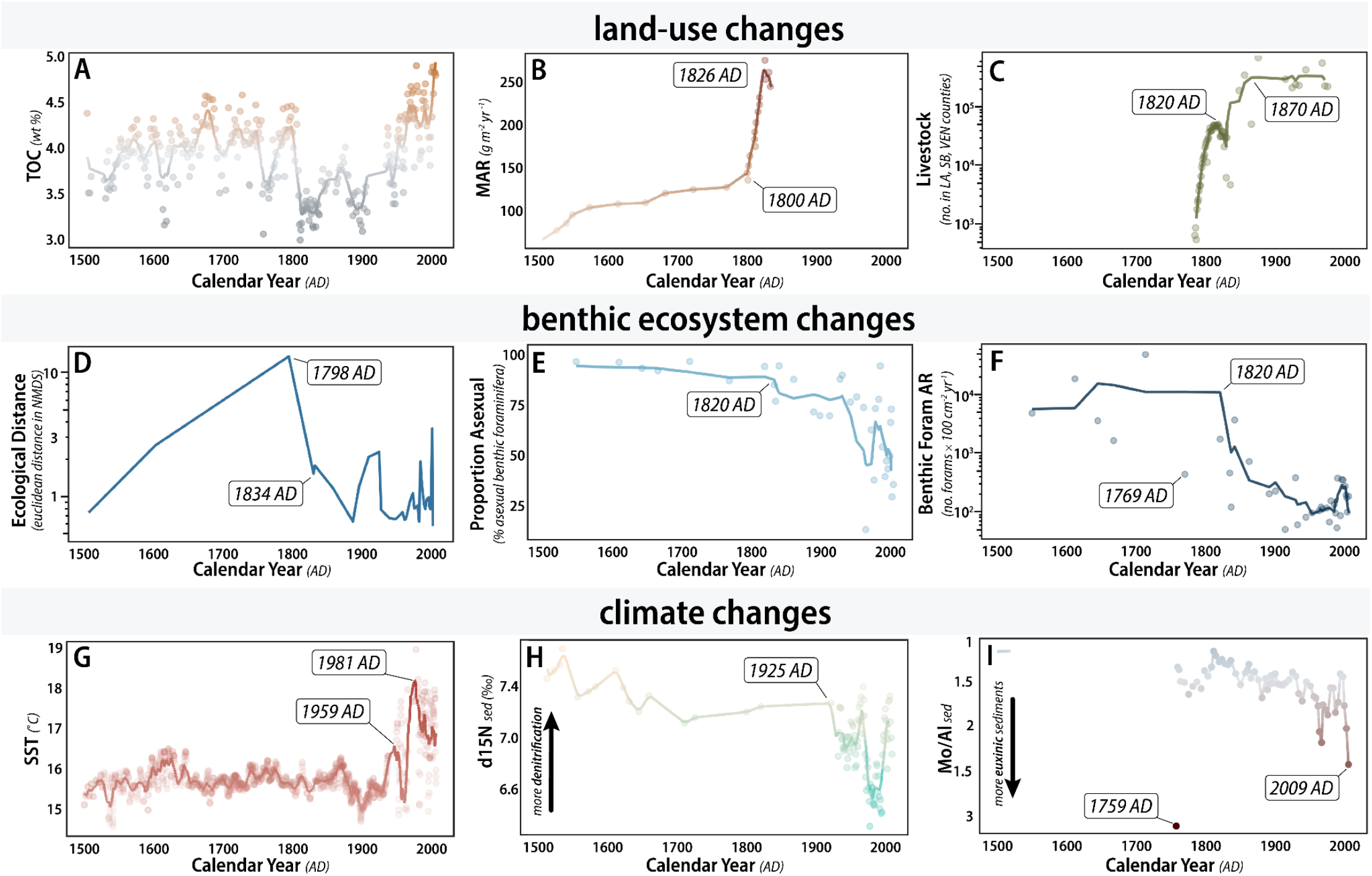
Proxies reflecting land-use, climate, and ecological changes in Southern California, 1500 AD to present. Panels (**A-C**): Proxy variables for land-use changes. (**A**) Total Organic Carbon (TOC; wt%) for SBB cores ODP 893 (*53*), SPR0901-03KC (*56*), and SBB-190629 MUC (*51*). (**B**) Mass Accumulation Rate (MAR; g m^-2^ yr^-1^) for SBB core SPR0901-06KC (*58*). (**C**) Total number of livestock (cows, horses, mules, sheep, pigs) for Los Angeles (LA), Santa Barbara (SB), and Ventura (VEN) counties, from California mission records (La Purisima Concepcion, Santa Barbara, Santa Ines) (*87*) and the U.S. Census of Agriculture. Panels (**D-F**)**: p**roxy variables for benthic ecosystem changes. (**D**) Ecological distance (euclidean distance in NMDS), this study, from cores MD02-2503 (*39*) and MV1012-BC1 (*48*). (**E**) Proportion asexual benthic foraminifera, an indicator of reproductive life history, from cores MV1012-KC1 and MV1012-BC1 (*80*). (**F**) Benthic foraminifer accumulation rate (number of foraminifera x 100 cm^-2^ yr^-1^), this study, from cores MV1012-KC1 and MV1012-BC1 (*48*). Panels (**G-I**): proxy variables for climate changes. (**G**) Sea Surface Temperature (SST; °C) estimated from alkenone (*49, 50*) and oxygen isotope proxy data (*41, 42*)and direct SST measurements from the CalCOFI monitoring program. (**H**) Sedimentary δ^15^N from SBB cores ODP 893 (*52*) and SBB-190629 MUC (*51*). Lower values of δ^15^N reflect more oxygenated source waters. **(I)** Sedimentary Mo/Al ratio from ODP 893 (*53*) and SPR0901-03KC (*54*). Higher values of Mo/Al reflect more euxinic sedimentary conditions (anoxia with free H_2_S). In each panel (except D), rolling 5-point averages are denoted by solid lines and points, while lighter points indicate values from original data.

Our work adds to growing evidence that colonial-era land-use change had an outsized ecological impact: we find that novel SBB ecosystem states already emerged in the early 1800s AD, beginning with the largest faunal turnover event of the past 34 kyr. We interpret the importance of TOC in predictive models as evidence for the role of terrestrial input in shaping the ecological composition and variability of benthic ecosystems on the California margin. While TOC is a complex oceanographic proxy (modulated by changes in surface ocean productivity, terrestrial deposition, and organic carbon preservation, among other factors), the majority (up to 80%) of TOC in the SBB is terrestrial in origin (*51, 81, 82*), and recent work suggests ∼15% of all organic carbon buried in the past 2 kyr originates from just 11 major flood events (*83*). Heightened TOC inputs to the SBB correlate with the marked increases in ecological distances we observe during the late Holocene (**Figure 2, Figure 3, SI Figure 8**).

Natural climate variation may have created the ideal background conditions for novel human land-use changes to drive the emergence of distinctly different ecosystem states in the 1800s AD. Ecological distance outliers in the 1360s and 1600s AD likely reflect benthic community responses to oceanographic conditions across the evolution of the Little Ice Age (LIA; **SI Figure 4, SI Figure 8**). Enhanced export productivity during early LIA cooling between 1200 and 1600 AD, (i.e., the transition from the Medieval Warm Period to the late LIA), is thought to have been driven by heightened oxygenation due to the changing influence of source waters to the CCS (*55*). However, CCS oxygen concentrations began to decline in the 1600s AD, potentially driven by sea ice expansion that reduced air-sea exchange (*55*). Beginning in the 1700s AD, spikes in sedimentary mass accumulation rates and secular increases in TOC due to land-use modification (**Figure 3; SI Figure 9**) may have further intensified bottom-water anoxia through the accelerated deposition of organic matter to the seafloor. Coming on the heels of the LIA, colonial-era land-use modification likely impacted benthic ecosystems just beginning to recover from widespread deoxygenation of the CCS.

Ultimately, the timing of the transition in the early 1800s to a novel ecosystem well outside the range of typical glacial-to-interglacial states is strong evidence that the anthropocene represents a fundamentally novel period in recent geologic history. The initial shift towards an anthropocene ecosystem state occurred too early to be explained by anthropogenically-forced climate warming, suggesting that colonial-era land-use change may have played a largely underrecognized role in driving early ecological regime shift in a deep-sea benthic ecosystem. This shift began decades to centuries before the advent of direct monitoring, with novel ecosystem states continuing through the present day–highlighting the importance of long-term paleontological data in understanding the structure of contemporary ecosystems. In addition, compositional similarities between communities from the late 2000s and the glacial-to-interglacial transition suggest that rapid warming and deoxygenation has consistent impacts on ecosystem structure across millennia, and that past changes may presage near-term impacts. Taken together, our work shows that humans have long had far-reaching impacts, even in the deep sea–and that the novel ecosystems predicted to emerge in the anthropocene (*21*) already arrived more than two centuries ago.

## Acknowledgments

We thank Jenni Brandon and William Jones for providing core samples from MV1012-BC1, and Tessa Hill and Hannah Palmer for constructive conversations during project conceptualization.

## Author contributions

Conceptualization: SSK, IAD, SF

Data Curation: SSK

Formal Analysis: SSK, SF, IAD, ILH, RDN

Funding acquisition: SSK, SF

Investigation: SSK, SF, IAD, ILH, RDN

Methodology: SSK, SF, IAD

Project Administration: SSK, SF

Resources: SF, RDN

Software: SSK, SF

Supervision: SSK, SF, IAD, ILH, RDN

Validation: SSK, SF, IAD, ILH, RDN

Visualization: SSK, SF, IAD

Writing – original draft: SSK, IAD, SF

Writing – review & editing: SSK, IAD, ILH, RDN, SF

## Data, code, and materials availability

All data and code used for this analysis are available in a permanent public repository accessible via the data citation in ref. (*84*)

## Supplementary Materials

Materials and Methods

Supplementary Text

Figs. S1 to S9

Tables S1 to S5

References (*85-137*)

Movie S1

