## Supplemental Materials for "Nineteenth-century emergence of a seafloor ecosystem beyond late Quaternary limits"

**The PDF file includes:**

Materials and Methods
Supplementary Text
Figs. S1 to S9
Tables S1 to S5
References

**Other Supplementary Materials for this manuscript include the following:**

Movie S1

### Materials and Methods

Sediment core collection

A box core from the center of the Santa Barbara Basin (SBB) was collected in 2010 at station MV1012-ST46.9 (34˚17.228'N, 120˚02.135'W, 580m water depth). To aid in chronology development, 2-cm vertical core slices were x-radiographed, line scanned at 1-mm intervals, and color photographed. An age model for core MV1012-BC1 was based on varve couplet dating and identification of stratigraphic marker events (*26*, *35*, *89*, *90*). Given its shallow depth, the traditional varve count ages assigned to MV1012-BC1 likely align well with a ΔR-corrected $^{14}C$ chronology developed by Hendy et al. (*26*). A regression model extended the chronology to the year 1834 of the Common Era (CE). Using this age model, the mean temporal grain for the box core was 2.2 yr/sample (minimum 1.14, maximum 2.7).

Benthic foraminifer community composition data

The box core was subsampled transversely every 0.5 cm, dried overnight at 50˚C, and washed with deionized water over 104- and 63-µm meshes. Due to their relatively small overall volumes, each of the 30 subsamples selected for this analysis was picked for all benthic foraminifera present under a Leica EZ4 dissecting microscope at 16x magnification. Picked benthic foraminifer assemblages were imaged using a Keyence VHX-7000 digital imaging microscope at 150x magnification, following high-throughput imaging techniques outlined in Kahanamoku et al. 2024. Full-assemblage images were segmented into individual images using the *AutoMorph* protocol, visually identified to species following references reported in Kahanamoku et al. 2024, and counted. Counts were transformed to accumulation rates (no. foraminifera x 100 $cm^{-2}$ $yr^{-1}$) to normalize for the temporal grain and surface area of the box core MV1012-BC1 (174 $cm^2$).

Data from a second core (MD02-2503) was obtained from Ohkushi et al. (*39*). Species names were updated for *Buliminella tenuata* to reflect the current accepted name, *Eubuliminella exilis*. Species counts were transformed to accumulation rates (no. foraminifera x 100 $cm^{-2}$ $yr^{-1}$) in order to normalize counts to the temporal grain and surface area of the jumbo piston core MD02-2503, reported by (*47*) as 95 $cm^2$.

A composite dataset was created by combining accumulation rates from cores MV1012-BC1 and MD02-2503. Species with a relative abundance of 30% or higher in any given sample were considered common; all others were grouped into a single category, "other," and accumulation rates were recalculated.

To account for the potential impacts of time-averaging, we ran all analyses reported in the main text and below on a composite dataset where counts from core MV1012 artificially time-averaged by combining original count data into ~7-year temporal bins to match the average resolution of core MD02-2503 (see "analytical approach to time averaging" below). All statistical analyses were conducted in R v. 4.4.1 (*91*). We used *vegan* package version 2.6.8 for R for ordinations (*92*).

NMDS and dissimilarity metrics

We performed nonmetric multidimensional scaling (a dimensionality reduction technique) on log-transformed species abundances, using euclidean distance as the dissimilarity metric (RMSE

= 0.003, stress = 0.066, non-metric fit $R^2 = 0.996$). Prior to log transformation, a small constant value (0.1, less than half the minimum non-zero species accumulation rate, chosen to be more conservative than the commonly-applied half-minimum convention) (*94*) was added to species accumulation rates to ensure that true absences remain distinguishable from rare occurrences with minimum distortion to the relative abundance structure of the data. We chose to use log transformation, rather than Hellinger transformation (a measure of relative abundance) in order to retain the influence of variability in species' accumulation rates. Accumulation rates are fluxes, representing time- and volume-normalized counts that partially account for variation in time-averaging (see below for further discussion on preservational considerations when interpreting accumulation rates). As a result, accumulation rates represent an ecologically-meaningful signal that captures changes to individual species' fluxes rather than their relative proportions. By using Euclidean dissimilarity on log-transformed accumulation rate data, our goal was to ensure that the role of species absences in determining community composition was taken into account in our ordination. While Bray-Curtis dissimilarity may be a more standard metric for ecological data, it ignores shared absences that are ecologically meaningful in our study system. In the Santa Barbara Basin, when analyzing time-averaged foraminiferal composition data, frequent species absences are expected to occur in response to minute variability in environmental drivers, such as hypoxia (*38*, *39*, *47*, *80*). As a result, species absences are suggestive of true absence within the SBB benthic foraminifer community and serve as a rich source of information on the environmental drivers of ecological change through time.

We grouped NMDS data into clusters using a k-means cluster analysis, where the optimal number of clusters (k = 3) was determined by computing the clustering algorithm for values of k = 1-10 and minimizing the within-cluster sum of squares (i.e., using the "elbow method"). As a secondary check, we calculated silhouette scores, which measure how similar an object is to its own cluster, and found k = 3 maximized cohesion (mean silhouette width = 0.5).

Environmental Proxy Data

To assess the influence of environmental variation on community composition, we used environmental proxy data generated from sediment cores sampled at or near the same locality as cores MD02-2503 and MV1012-BC1 (**SI Table 1**).

*Sea Surface Temperature*

To create a 34-kyr-long record of sea surface temperature (SST, ˚C), we combined SST estimates from three primary sources: $^{16}O/^{18}O$ ratios ($\delta^{18}O$, ‰) reported in (*42*) for cores MV0508-11JPC, -16JPC, and -20JPC from 59.9 ka to 1592 AD; the alkenone $U^{k'}_{37}$ index reported in (*49*) for box cores SABA87-1, SABA87-2, and SABA88-1 from 1440 AD to 1940 AD; and direct SST measurements from the California Cooperative Oceanic Fisheries Investigations (CalCOFI) program from 1953 AD to the top of our core record in 2008 AD (https://calcofi.org).

Specifically, we used an estimate of SST reported by (*42*) based on a time series of $\delta^{18}O$ from the planktonic foraminifer *Globigerinoides bulloides*, corrected using a species-specific temperature calibration and accounting for ice-volume effects and modern local salinity. They estimate temperature as:

$$T\ (^{\circ}C) = 14.62 - 4.70(\delta^{18}O_c - \delta^{18}O_w)$$

Where $\delta^{18}Oc$ represents the $\delta^{18}O$ of foraminiferal calcite and $\delta^{18}Ow$ represents the $\delta^{18}O$ of seawater. This equation is calibrated from 2˚C to 25˚C with an error of 1.24 ˚C (1σ) (*95*). *G. bulloides* is a common planktonic foraminifer species that is considered to primarily inhabit cool upwelled waters (*96–98*), and is associated with upwelling in the Southern California bight (*99*, *100*).

The alkenone $U^{k'}_{37}$ index reflects the temperature-dependent unsaturation of long-chain ($C_{37}$) ketone lipids (alkenones) produced by haptophyte algae (*101–103*) and well-preserved in ocean sediments. Alkenones are produced in the highest amounts when haptophyte populations are large (*104*, *105*), such that $U^{k'}_{37}$ values are thought to reflect the seasonality of haptophyte blooms (*106*, *107*).

We included direct SST measurements from the 70-year-long records of the California Cooperative Oceanic Fisheries Investigations (CalCOFI) from CalCOFI Station 81.8/46.9 (34'16.49˚N, 120'1.51˚W), located in the center of the Santa Barbara Basin and directly overlying the locality for sediment cores MV1012-BC1 and MD02-2503. To ensure that direct SST measurements were comparable to proxy estimates, we selected data from the CalCOFI bottle database (1953 AD to present; https://calcofi.org/data/oceanographic-data/bottle-database/) that reflected SST measurements taken during upwelling season when alkenone production should be most pronounced. We considered samples from Quarters 2 and 3 (April-June, July-September) to be most reflective of intervals of heightened upwelling in the CCS, which typically occurs between April and September (*108*, *109*), and estimated median, mean, and maximum temperatures for each year between 1946 AD and 2008 AD. Comparing summary statistics for upwelling-season direct SST measurements and alkenone SST estimates showed that alkenones appeared to best correlate to maximum temperatures. As a result, our analyses incorporate maximum SST for the 1953-2008 AD portion of our record.

All three variable types ( $\delta^{18}O$, alkenone $U^{k'}_{37}$ index, and upwelling-season maximum SST from CalCOFI) were combined into a single record and missing years were interpolated linearly.

*Nitrogen isotopes ($\delta^{15}N$)*

The $^{15}N/^{14}N$ ratio of sediments ($\delta^{15}N$; ‰) is modulated by the production of nitrogen-bearing organic matter in the surface ocean and its burial in the sediments. As a result, $\delta^{15}N$ reflects aspects of the marine nitrogen cycle in both the water column and within sediments. In the California Current System, $\delta^{15}N$ primarily records the intensity of denitrification in the oxygen-poor Eastern Tropical North Pacific (*72*, *110–112*), modified by the variable influence of well-oxygenated North Pacific Intermediate Waters (*56*, *113–115*).

We sourced data on $\delta^{15}N$ from two publications: Emmer and Thunell (*52*) for core ODP 893 from 48.6 ka to 1927 AD, and Xu et al. (*51*) for core SBB-190629 MUC from 1934 AD to 2018 AD. To fill gaps in the data from ODP 893, we used linear interpolation to estimate values from missing years.

*Total Organic Carbon*

Total Organic Carbon (TOC, wt%) is often used as a proxy for carbon export from the upper ocean. However, the majority (up to 80%) of TOC in SBB sediments is terrestrial in origin (*51*,

*82*, *116*). Large terrestrial depositional events ("megafloods") are primary sources of organic carbon burial, with a recent study estimating that 15% of all organic carbon buried in the SBB over the past 2 kyr originated from just 11 major flood events (*83*). Flood events are a considerable source of terrestrial organic carbon to the California margin, and may act to further enhance TOC values through their impact on sedimentary preservation and/or diagenesis. Higher TOC values, likely resulting from enhanced organic carbon preservation, may occur beneath instantaneous depositional events (e.g., both floods and turbidites) as rapid mobilization of sediments is thought to reduce $O_2$ penetration and aerobic organic carbon degradation (*56*).

For this analysis, data on TOC was derived from three sources: Ivanochko and Pedersen (*53*) for core ODP 893 from 48.9 ka to 0.357 ka, Wang et al. (*56*) for core SPR0901-03KC from 2.1 ka to 1910 AD, and Xu et al. (*51*) for core SBB-190629 MUC from 1934 AD to 2018 AD. We used linear interpolation to estimate sedimentary TOC values for years where records do not overlap.

<u>*Sedimentary Redox*</u>

The oxygenation state of pore waters within sediments drives changes to sedimentary redox conditions and influences the mobility and accumulation of metals within sediment. As deoxygenation develops, organisms utilize secondary oxidant sources for anaerobic metabolic processes (e.g., respiration, organic matter degradation). Redox reactions are classically considered to progress as follows: (1) denitrification, (2) manganese reduction, (3) iron reduction, and (4) sulfate reduction (*117*). Under reducing conditions, certain trace metals within sediments enrich to levels well above crustal conditions. In paleoceanography, enrichment of the trace metals Molybdenum (Mo) and Rhenium (Re) is often used as a proxy for past redox states. Each element records different aspects of redox cycles: Re enrichment is thought to indicate suboxic (absence of $O_2$ and $H_2S$) conditions, while Mo enrichment indicates euxinic (absence of $O_2$ and free $H_2S$) conditions (*73*, *118*). Because variability in the composition of sediments through time can dilute abundance of trace elements, trace-element concentrations are often normalized to aluminum content (*119*).

Trace-metal proxies for sedimentary redox conditions were derived from two sources: Ivanochko and Pedersen (*53*) for ODP 893 from 48.5 ka to 1807 AD, and Wang et al. (*54*) for core SPR0901-04BC from 1759 AD to 2008 AD.

<u>*Sedimentary Mass Accumulation Rate*</u>

Sedimentary Mass Accumulation Rate (MAR; g $cm^{-2}$ $yr^{-1}$) is a measure of sediment deposition over time that takes into account the mass of sediment deposited per unit area alongside changes to sediment porosity. For this analysis, data on sedimentary MAR was derived from three sources: Nederbragt et al. (*57*) which reports data for core ODP 893 from 16 to 0 ka; Du et al. (*58*), which reports data for cores MV0811-14JC and SPR0901-06KC from 9000 BCE to 1834 CE, and Xu et al. (*51*), which reports data MAR for core SBB-190629 MUC from 1937-2008 CE. Du et al. and Xu et al. directly reported sedimentary MAR in g $m^{-2}$ $yr^{-1}$. We calculated sedimentary MAR for ODP 893 using porosity and dry sediment weights reported in Nederbracht et al. 2008 and porosity data reported in Kennett et al. 1994 following equations 4 and 5 in Xu et al. (*51*):

$DBD = p_s\ x\ (1\text{-}\varphi)$, where $p_s$ is sediment bulk density (g $cm^{-3}$) and φ is porosity; and

$MAR = LSR \times DBD \times 10{,}000$, where LSR is linear sedimentation rate (cm $yr^{-1}$)

We used linear interpolation to estimate sedimentary MAR values for samples between 1834 and 1937 AD where records do not overlap.

### Supplementary Text

In the Santa Barbara Basin (SBB) of Southern California, persistent hypoxia has shaped a benthic ecosystem composed primarily of low-oxygen specialists. Benthic foraminifera represent the majority of eukaryotic biomass from the central SBB (*36*, *37*), as these unicellular, shelled eukaryotes are are uniquely able to survive persistent hypoxia due to the variety of metabolic adaptations that allow them to respire both aerobically and anaerobically (*66*, *120–122*). Intraspecific differences in hypoxia tolerance makes foraminifer assemblages sensitive recorders of minute changes in dissolved oxygen concentrations through time. Given their high fossilization potential, benthic foraminifera are a key tool in the reconstruction of past climate and oceanographic conditions. As a result, there is rich data available for these taxa spanning the last glacial period and through several intervals of rapid and extreme warming in the SBB (*34*, *38*, *39*, *47*, *123*).

The California Current System (CCS) is among the few marine systems where direct observations can be coupled with seasonal-to-annual paleontological and paleoclimate data (Schimmelman et al. 1996) to retroactively extend ecosystem monitoring prior to the onset of anthropogenic climate change. Seasonal upwelling of nutrient-rich waters within the CCS drives deep-water deoxygenation via enhanced export production of organic matter from the photic zone, facilitating the formation of millimeter-scale seasonal varve couplets within the persistently hypoxic borderland basins of Southern California (Reimers et al. 1990) due to an almost complete lack of bioturbation. Sedimentary archives from these sites record interannual climate, hydrographic, and biological variability within the broader CCS over the past tens of thousands of years (*29*, *30*, *33*, *59*). This extensive past data record, along with several long-term monitoring programs in operation since the 1950s (including the California Cooperative Oceanic Fisheries Investigations, or CalCOFI; https://calcofi.org), makes the CCS one of the best-observed marine ecosystem complexes in the world (*31*, *124*).

*A note on terminology*

Throughout this publication, we use the term 'ecosystem' rather than 'community' deliberately. Because no study can fully characterize an ecosystem in the complete conceptual sense, the appropriateness of the term is a matter of degree rather than a binary judgment. We argue that 'ecosystem' is the more fitting choice here for two reasons. First, our focus is on the complex interactions between biotic responses and abiotic forcing—including links between photic zone processes, deep-sea benthic dynamics, and land-sea interactions—and framing these as mere 'community' changes would not capture the systems-level nature of our findings. Second, given the unusual characteristics of the SBB benthic environment, a study centered on foraminifera against a backdrop of key abiotic parameters (land-use change, ocean warming and deoxygenation, etc.) likely captures a comparably high proportion of true ecosystem-level variance relative to what is typically achieved in studies of more speciose and less tractable ecosystems.

We also acknowledge that the term anthropocene carries several connotations. In geology, the term “Anthropocene” is likely considered to equate to the proposed, unofficial, and recently rejected geologic epoch of the same name. We agree with the recent decision by the International Commission on Stratigraphy to reject the Anthropocene as a formal unit on the geologic timescale (*13*), as the mismatches in temporal resolution between proposed chronostratigraphic markers for the beginning of an “Anthropocene Epoch” make it exceedingly difficult to assign a global, uniform chronostratigraphic boundary . However, we recognize the utility of the term “anthropocene” as an acknowledgement of the magnitude, variety, and longevity of human-induced changes on Earth's ecosystems and climate (*12*). We consider our findings of an early 1800s regime shift towards an anthropocene-type ecosystem to be reflective of a broader interpretation of the “anthropocene”as the period over which the “intensified repetition of anthropogenic environmental change [beginning with] colonial practices that facilitated [global] industrial expansion” (*125*) (p. 156) (see also refs (*126*, *127*)). Thus, our use of the term anthropocene (decapitalized) seeks to capture the anthropocene *sensu lato*–the past several centuries over which novel human impacts had outsize effects on global ecosystems on land and in the oceans.

<u>Faunal categories</u>

NMDS cluster groupings are reported in **SI Table 2**. NMDS species vectors broadly reflect hypoxia tolerance (**SI Figure 2**). Interpretations of species’ environmental preferences were based on known oxygen tolerances (*128–131*) and inferred trophic habitats (*122*, *132*, *133*). NMDS species groupings are as follows: **Group 1 (Hypoxia-Tolerant Detritivores):** *Suggrunda eckisi, Fursenkoina cornuta,* and *Chilostomella ovoidea*; **Group 2 (Hypoxia-tolerant denitrifiers):** *Bolivina advena, Bolivina argentea, Buliminella curta, Bolivina pseudobeyrichi, Bolivina interjuncta, Bolivina spissa, Eubuliminella exilis,* and *Nonionella stella*; **Group 3 (Ephemeral hypoxia tolerance):** *Anglogerina carinata, Bolivina tumida, Cassidulina limbata, Globobulimina auriculata, Globobulimina pacifica,* and *Uvigerina perigrina*; **Group 4 (Oxic taxa):** *Cassidulina translucens, Epistominella pacifica, Epistominella smithi, Fursenkoina rotunda, Nonionellina labradorica, Quinqueloculina seminulum,* and *Quinqueloculina spp.* **Group 5 (Modern OMZ):** *Bolivina spissa, Uvigerina peregrina,* and all other rare taxa; (**Glacial Taxa):** *Epistominella pacifica, Epistominella smithi, Globobulimina pacifica, Nonionellina labradoria, Quinqueloculina seminulum,* and *Quinqueloculina* spp.

<u>Faunal die-offs and extremophile abundance through warming events</u>
Complete faunal die-offs occurred during the Bolling warming (14.444 ka) and the pre-Boreal warming (between 11.633 ka-11.626 ka). These die-off events have been previously interpreted as reflecting intervals of prolonged total anoxia in the deep SBB (Ohkushi et al. 2013). Extreme deoxygenation likely lasted for several decades, evidenced by low abundances of individual foraminifera in several samples surrounding die-off events (e.g., <10 from 11.648 and 11.596 ka). Sharp increases in the relative abundance of *Nonionella stella* just prior to or following die-off events provide additional evidence for anoxic conditions. *N. stella* is considered an “anoxic

extremophile" that can survive for several months in total anoxia (0 ml $L^{-1}$ $O_2$) as well as under euxinic conditions (total anoxia combined with the presence of free $H_2S$) (*36*, *64–66*, *134*, *135*). *N. stella* reached peak relative abundances during three intervals of extreme deoxygenation: during the Bolling warming just prior to the die-off event at 14.524 ka (50% relative abundance) and during the pre-Boreal just following the die-off interval (15%).

Ecological distance over time

We calculated ecological distance as euclidean distances between consecutive benthic foraminiferal assemblages. The Euclidean distance matrix underpinning NMDS results was used for this analysis. We considered outliers for ecological distances to be representative of large, abrupt faunal turnover events (i.e., saltatory change). To compare anthropocene-group data to a historical baseline, we calculated outliers as above 1.96σ for the pooled Euclidean distances of samples older than 1 ka. Ecological distance outliers are reported in **SI Table 3**. We also calculated a historical maximum in Euclidean distance by finding the maximum value for all data older than 1 ka. **SI Figure 4** provides an overview of these results.

Analytical approach to time-averaging

Temporal mixing of sedimentological deposits can result in time-averaging, where individual strata may contain fossils and sedimentary structures that pre- or post-date its deposition. Time-averaging can mask true paleontological trends by asynchronous events appear synchronous in the geologic record. Our analysis may be susceptible to the impacts of time-averaging due to the variable resolution between youngest and oldest samples (~1 yr/sample at the coretop of MV1012-BC1, ~4 yr/sample at the bottom of MV1012-BC1, and ~7 yr/sample on average in MD02-2503). As a result, it is possible that heightened variability in the Anthropocene cluster group (which includes samples from both cores spanning 1-7 yr/sample resolution) compared to the older samples in the interglacial and glacial groups is an artifact of the higher resolution present in the youngest samples from our time series.

We addressed time-averaging issues analytically by binning samples from core MV1012-BC1 into ~7-year time bins and re-running NMDS ordinations, clustering, and ecological distance (euclidean distance in NMDS) using these binned data. We find that results from un-binned and binned data are similar (**SI Figure 5**).

Environmental correlates of community composition and ecological distance

We used generalized linear models (GLMs) and linear mixed models (LMMs), and a gradient boosting classification machine (GBM) to evaluate environmental controls on community composition and ecological distances through time. While GLMs and LMMS, are well-suited to detecting linear or smoothly-varying relationships between environmental predictors and biological response variables, ecological systems often encompass multiple environmental drivers interacting in complex, non-linear ways that these models may not fully capture. A GBM addresses this with an ensemble approach to approximate complex interactions among predictors. Importantly, this process does not require relationships to be specified in advance, and it produces a direct ranking of each predictor's contribution to explaining variability in the response—analogous to, but independent of, the $R^2$ values produced by GLM/LMMs.

34-kyr trends for environmental proxy variables used in regression models are shown in **SI Figure 6**. We compared several linear models examining the relationship between environmental proxy variables and MDS axis 1. A model with TOC, Mo/Al, and $\delta^{15}N$ as predictors of MDS1 was the best fit (**SI Table 4**). In single-predictor models, TOC was the best predictor of NMDS outputs ($p < 0.001$, $R^2 = 0.74$), followed by $\delta^{15}N$ ($p < 0.001$, $R^2 = 0.51$) and Mo/Al ($p < 0.001$, $R^2 = 0.34$). A GBM reflects this relationship (**Figure 2A**), where TOC explains the highest proportion of variance in community composition (gain = 68%). In a linear mixed model including TOC, Mo/Al, $\delta^{15}N$ and cluster group as a random effect, TOC and $\delta^{15}N$ were significant predictors of MDS1 but had a low marginal $R^2$ (0.05; **SI Table 4**). A visualization of the relationships between significant proxy variables and MDS axes 1 and 2 can be found in **SI Figure 7**.

We compared several linear models examining the relationship between environmental proxy variables and ecological distance. Two models had a statistically identical best fit: one with TOC as the only predictor of ecological distance, and another including both TOC and Mo/Al (**SI Table 5**). In a linear mixed model including TOC, Mo/Al, $\delta^{15}N$ and cluster group as a random effect had a relatively low marginal $R^2$ (0.14; **SI Table 5**). Relationships between significant proxy variables and ecological distance can be visualized in **SI Figure 8**.

<u>$\delta^{15}N$ as an indicator of ETNP denitrification</u>

An important consideration in the interpretation of sedimentary $\delta^{15}N$ records from the SBB is the potential contribution of benthic denitrification to isotopic enrichment. Given that the bulk of benthic biomass in the SBB consists of prolific denitrifiers (including benthic foraminifera), it is important to consider whether sedimentary denitrification meaningfully modifies the $\delta^{15}N$ signal independent of water column processes. This question was directly addressed by Sigman et al. (*136*), who sought to explain why the degree of isotopic enrichment in SBB sediments was substantially lower than expected given the large nitrate deficit in the basin. They demonstrated that despite high rates of sedimentary denitrification in the SBB, overall isotopic enrichment remains low and comparable to ETNP values, potentially as a result of high rates of nitrate resupply within the sediments that effectively buffer the benthic denitrification signal. These findings support the use of sedimentary $\delta^{15}N$ as a reliable recorder of source-water denitrification in the SBB, rather than a mixed signal confounded by benthic processes.

<u>TOC as an indicator of terrestrial input and preservational considerations</u>

Total organic carbon (TOC) in SBB sediments shows significant variability on both multi-millennial and multi-centennial timescales (**SI Figure 6**, **Main Text Figure 3**). Over the past 2 kyr, increases in TOC concentrations of ~1 wt% are generally interpreted as reflecting increased export productivity (*56*) (Wang et al. 2019). However, TOC records from the past 500 years show minimum values in the 1800s AD that persisted through the early 1900s, which we interpret as potentially reflecting increased terrestrial input resulting from land-use change in the source watersheds of the SBB—specifically the Santa Ynez, Santa Clara, and Ventura watersheds (*76*, *82*). This interpretation is supported by the observation that TOC decreases by ~1.5 wt% during flood events relative to background conditions (*83*), consistent with dilution by refractory terrestrial organic matter. An important caveat to the use of TOC as a proxy for export productivity is that preserved TOC may overrepresent the terrestrial fraction of original organic matter input due to taphonomic effects. Specifically, terrestrial organic matter is rich in recalcitrant compounds (e.g., lignin), whereas organic matter derived from local export production is comparatively labile and subject to much higher rates of decomposition prior to burial (*137*). As a result, preserved TOC concentrations may not accurately reflect the relative proportions of terrestrial versus photic zone particulate organic carbon originally supplied to the benthic environment, and the nutritional and redox impacts of these two fractions on the benthic ecosystem are likely to differ substantially.

In the 1800s AD, several event layers represent an additional source of complexity in interpreting the influence of TOC on benthic foraminiferal assemblages. First, an earthquake dated to 1812 AD and associated with a sedimentary event layer would have subjected benthic foraminifera to rapid burial. This event was closely followed by a turbidite exceeding 5 cm in thickness associated with the '*Macoma'* event' (when SBB bottom-water oxygen concentrations were high enough to support the brief flourishing of a macrofaunal community between ~1835 and 1840 AD) (*35*). Both events introduce discontinuities in proxy records from the SBB and complicate the interpretation of assemblage changes in their immediate aftermath.

Interpreting species accumulation rates and taphonomic considerations

Species' accumulation rates are indirectly related to abundance in an ecological sense, as they represent the number of dead foraminifer tests (i.e., shells) integrated over time and are thus dependent on changes to biological production, as well as abiotic factors that influence deposition and preservation post-mortem. Foraminifer production is modulated by life history factors, including reproductive frequency and output, lifespan, and mortality rate. The preservation of foraminiferal tests can also affect inferred accumulation rates (e.g., if tests are removed via dissolution or disintegration or altered by preservational processes in a way that obstructs identification).

However, in the Santa Barbara Basin, preservational impacts are considered to be minimal; persistent hypoxia limits vertical migration in sediments and preserves mm-scale structures within lamina, including submillimeter microhabitat structure within sediments (*133*). Calcium-carbonate foraminifer tests have high preservation potential, and, in the SBB, are typically well-preserved with no visual evidence of dissolution (*48*). In addition, previous work in the SBB demonstrates that the infaunal range of living foraminifera is limited to the first 1 cm of sediment (or significantly less, e.g., 1 mm for hypoxia-tolerant but anoxia-intolerant such as *Bolivina alata, Bulimina exilis*, and *Fursenkoina cornuta*) (*133*). As a result, foraminifer accumulation rates from the SBB are generally considered to primarily reflect changes to the production of foraminifer tests, rather than the influence of post-depositional processes.

Land-use, climate, and ecosystem changes in the SBB through time

Previous work in the CCS suggests that changes to terrestrial input to the deep continental margin may have driven rapid changes to ecological structure and stability through its compounding effects on oxygen loss even during the deglacial transition. Hendy and Pedersen (*113*) found evidence for large temporal changes in export production and variability in intermediate water oxygen supply in the CCS. During the Bølling interval (~14.5 ka) when near-total anoxia may have caused decades-long die-offs of deep benthic ecosystems in the SBB, rapid sedimentation alongside high rates of export production may have driven near-surface sediments towards strongly reducing conditions, reflected in concurrent peaks in Mo enrichment (**SI Figure 5**).

Over the past several centuries, concurrent changes to terrestrial input and oxygenation occur across an interval with compounding climatological and human stressors. Benthic ecosystem changes beginning in the early 1800s AD correspond with the growth of livestock populations in Southern California and resulting increases in sediment load to the Southern California Borderland basins (**SI Figure 9**). These changes predate the impacts from exponential population growth in the 20$^{th}$ and 21$^{st}$ centuries, as well as local expressions of anthropogenically-forced climate warming within the California Current System (**Main Text Figure 3**).

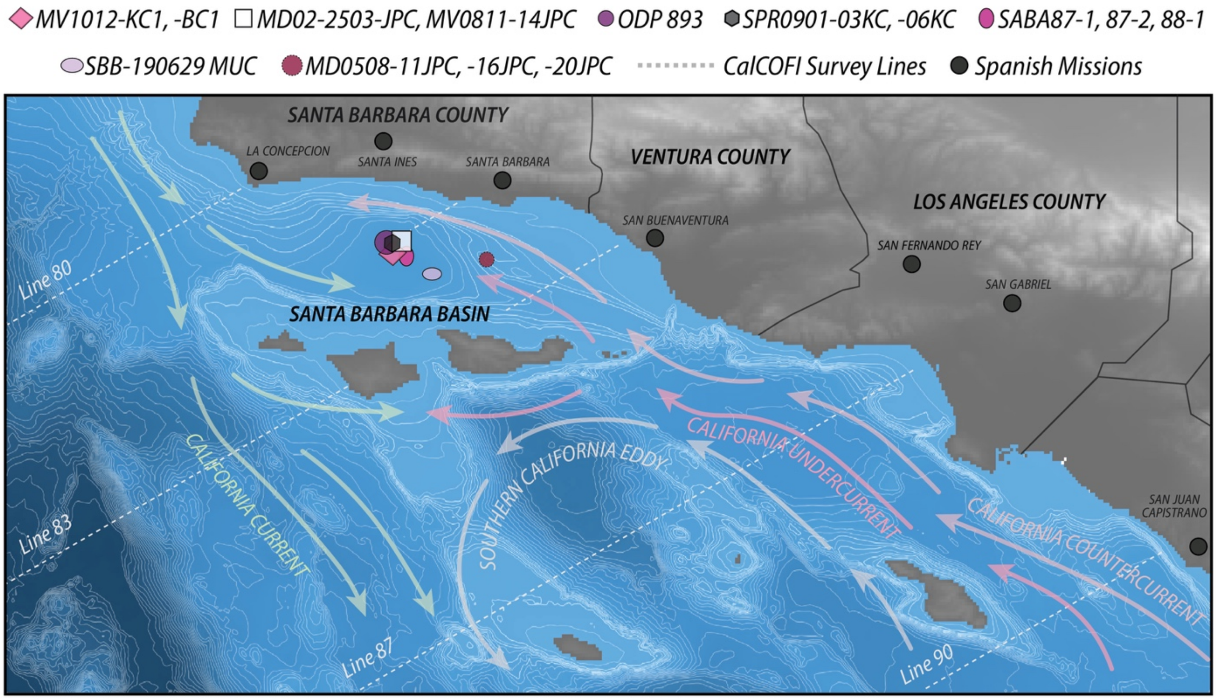


**Fig. S1. Map of Santa Barbara Basin core localities in the Southern California Bight.** Points show location of cores ODP 893, MD02-2503, MV1012-KC1, MV1012-BC1, SPR0901-03KC, SPR0901-06KC, and SBB-190629 MUC within central Santa Barbara Basin. Dotted lines indicate CalCOFI survey sites; survey line numbers are labeled. General direction of the California Current, California Countercurrent (inshore), California Undercurrent (at depth), and Southern California Eddy denoted by colored arrows. Black circles show locations of nearby Southern California missions, in order from north to south: La Purísima Concepción, Santa Inés, Santa Barbara, San Buenaventura, San Fernando Rey, San Gabriel, and San Juan Capistrano. Shading corresponds to GEBCO bathymetry (light blue: shallow; dark blue: deep) with 50-m contour lines overlain in white. Alpha values denote elevation (dark: lower elevation; light: higher elevation). DEM information from USGS (*93*).

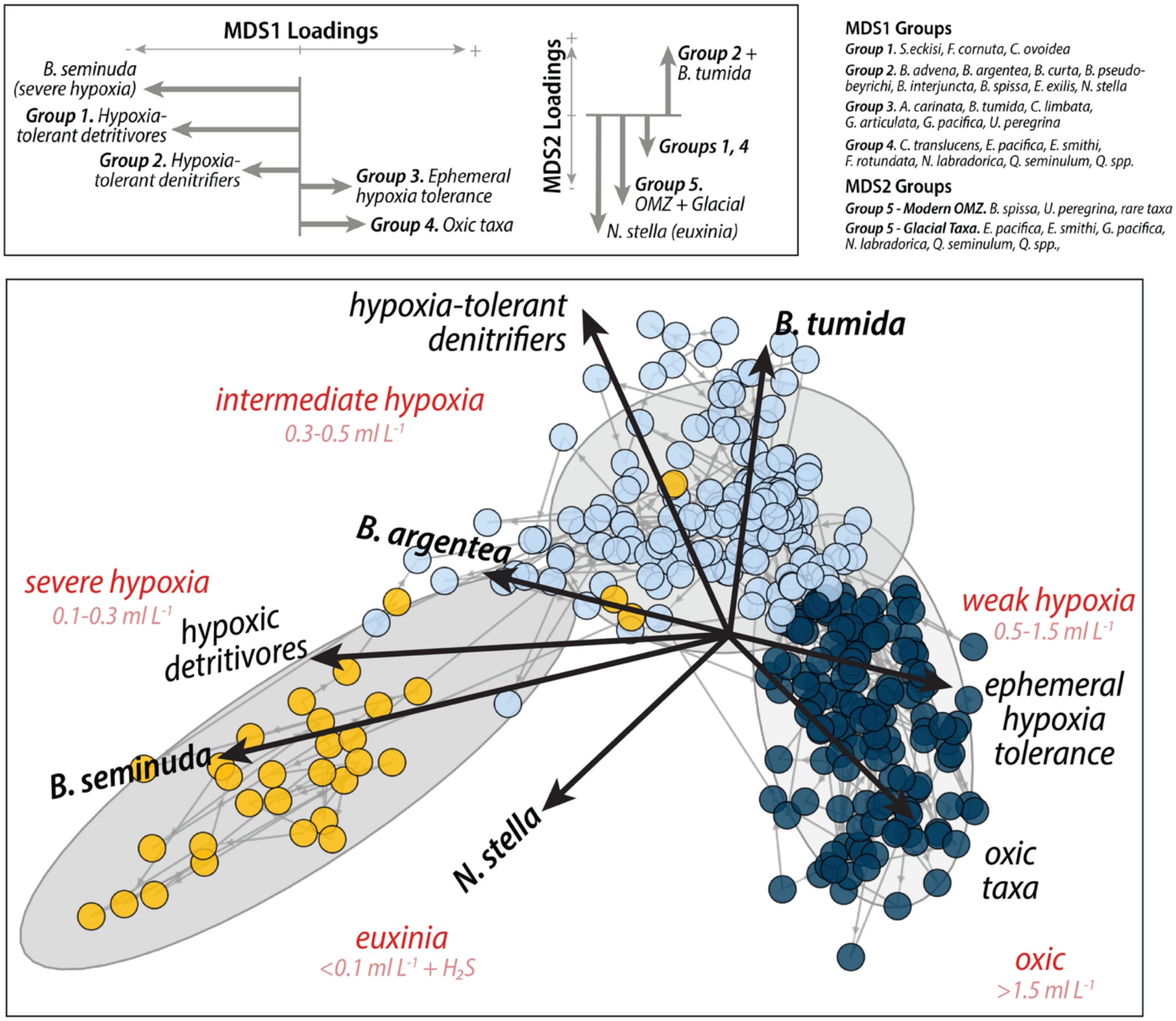


**Fig. S2. Non-metric multidimensional scaling (NMDS) ordination of benthic foraminiferal assemblages with species vectors and oxygen interpretations.**

Points are colored by cluster group: dark blue = glacial stadial, light blue = interglacial and glacial interstadial, and gold = anthropocene. Grey ellipses denote k-means clusters. Arrows show species vectors alongside oxygen tolerances and inferred trophic habits of primary species groups. Oxygen affinities represent the following DO concentrations: oxic (>1.5ml $L^{-1}$ $O_2$), weakly hypoxic (0.5-1.5 ml $L^{-1}$ $O_2$), intermediate hypoxia (0.3-0.5 ml $L^{-1}$ $O_2$), severe hypoxia (0.1-03 ml $L^{-1}$ $O_2$) and euxinia (<0.1 ml $L^{-1}$ $O_2$ + free $H_2S$). Top insets show species groupings and inferred trophic habitats and oxygen tolerances on MDS axes 1 and 2.

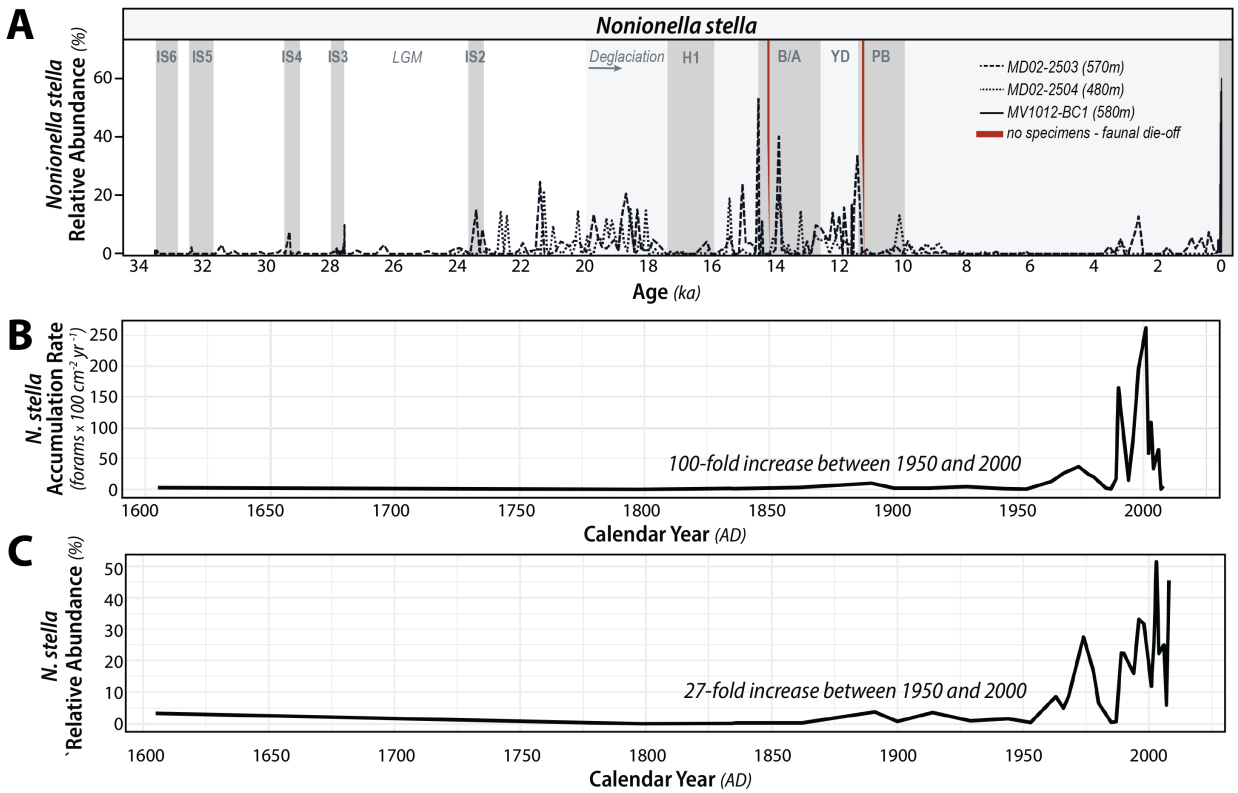


**Fig. S3.** ***Nonionella stella*** **relative abundances and accumulation rates through time.**
**(A)** *N. stella* relative abundance, 34 kya to present, from cores MD02-2503 and MV1012-BC1 (data used in main analysis) as well as a shallower core, MD02-2504 (*34*) to illustrate assemblage trends at shallower sill depths in the SBB (i.e., outside of the long-term anoxic core). Grey bars denote interstadial events (IS: interstadials 6 through 2; H1: Heinrich event 1; B/A: Bolling and Allerod warming events; YD: Younger Dryas; PB: pre-boreal warming. Red bars denote intervals during which no specimens were found in core samples, interpreted as complete faunal die-off events. **(B-C):** *N. stella* accumulation rate **(B)** and relative abundance **(C)** between 1600 AD and 2008 AD (the top of core MV1012-BC1).

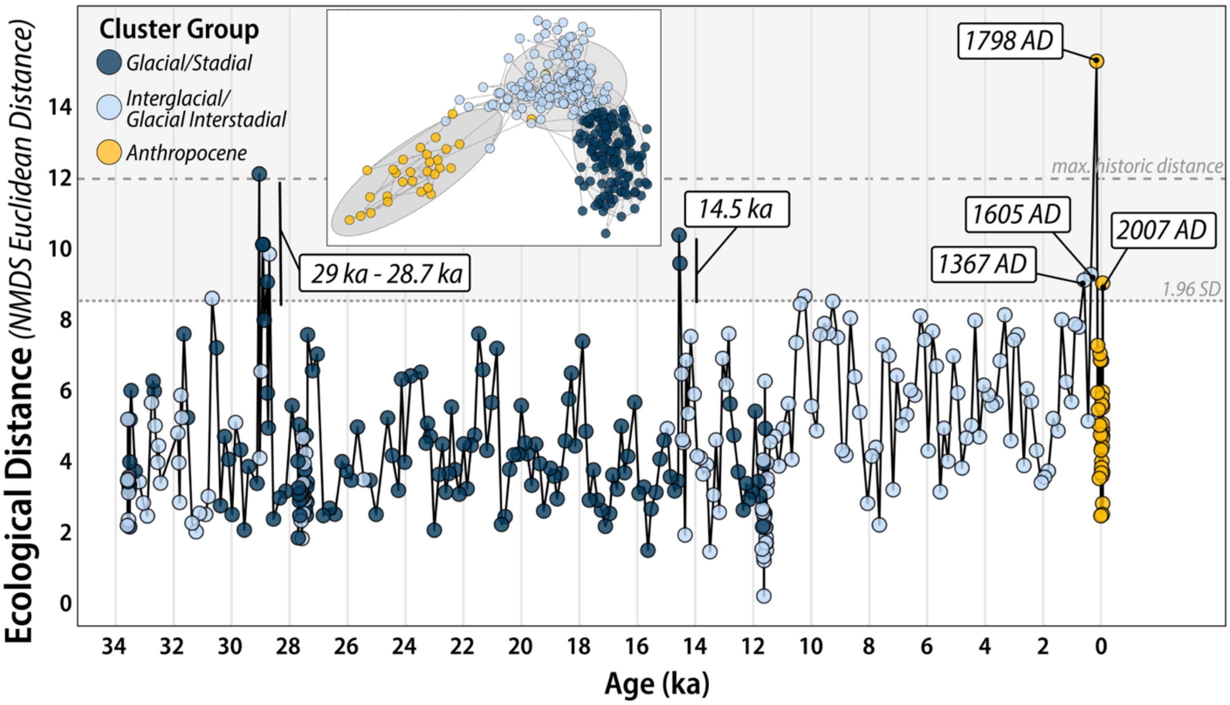


**Fig. S4. Ecological distance over time.**

Ecological distance is represented here as euclidean distance between foraminiferal assemblages. Colors denote k-means cluster grouping and age (dark blue: Glacial/Stadial, light blue: Interglacial/Interstadial; yellow: anthropocene). Dashed and dotted lines denote 1.96 SD and maximum historic value for ecological distance prior to 1798 AD. Points at or above 1.96 SD are labeled with age (ka) or calendar year (AD).

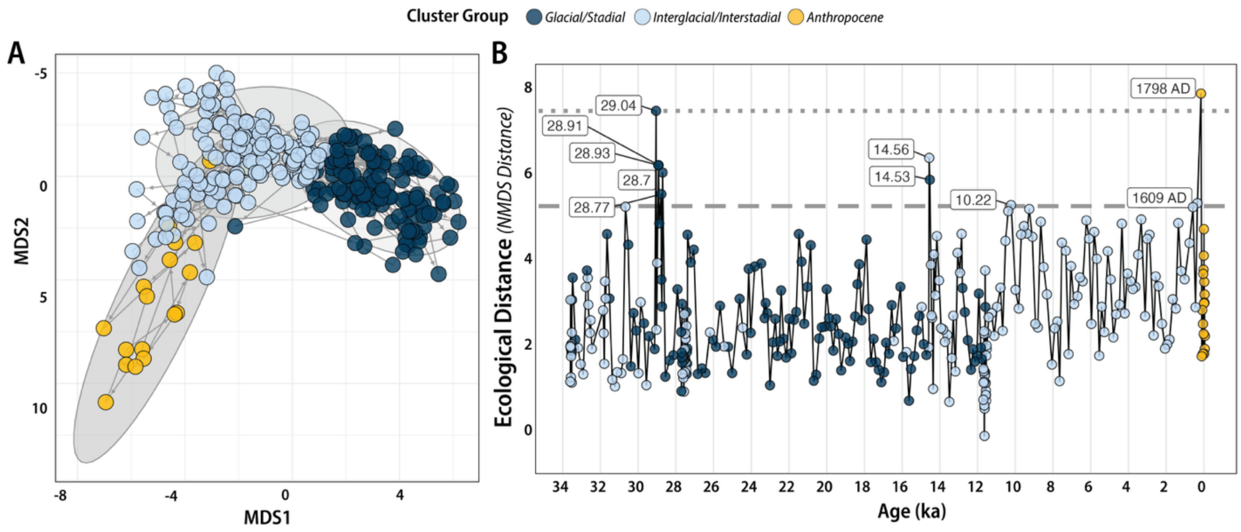


**Fig. S5. Artificially time-averaged data produce similar results.**

(**A**) NMDS and (**B**) Ecological Distance, recalculated using artificially time-averaged data (~7-year bins assigned to core MV1012-BC1 and coretop of MD02-2503 to match the average temporal resolution of core MD02-2503).

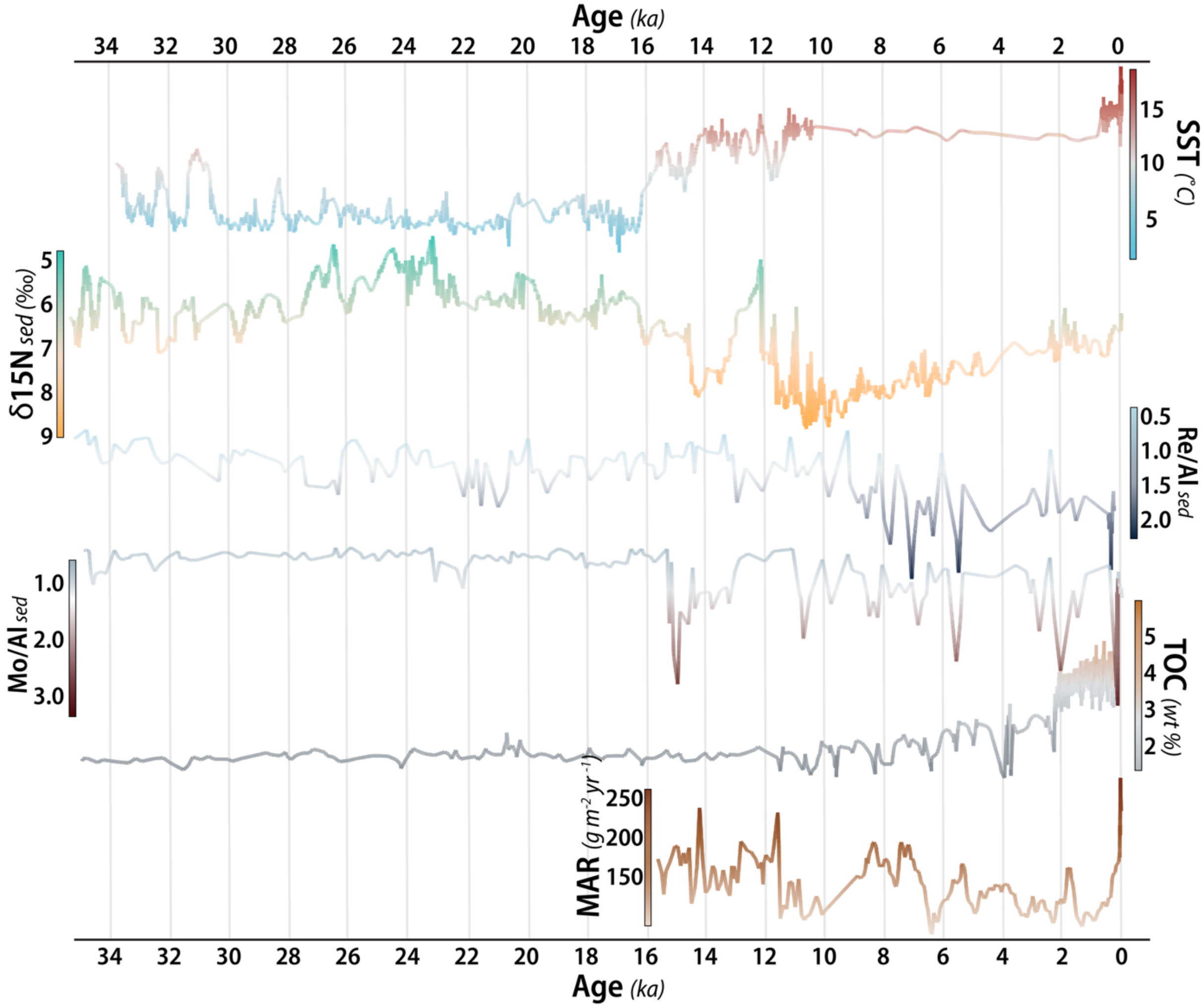


**Fig. S6. Environmental trends in the Southern California Bight, 34 ka to present.**

From top to bottom: sea surface temperature (SST, ˚C); sedimentary nitrogen isotopes ($\delta^{15}N$, ‰); sedimentary rhenium enrichment (normalized to aluminum, as Re/Al); sedimentary molybdenum enrichment (normalized to aluminum, Mo/Al); total organic carbon (TOC, wt %); and sedimentary mass accumulation rate (MAR, g $m^{-2}$ $yr^{-1}$).

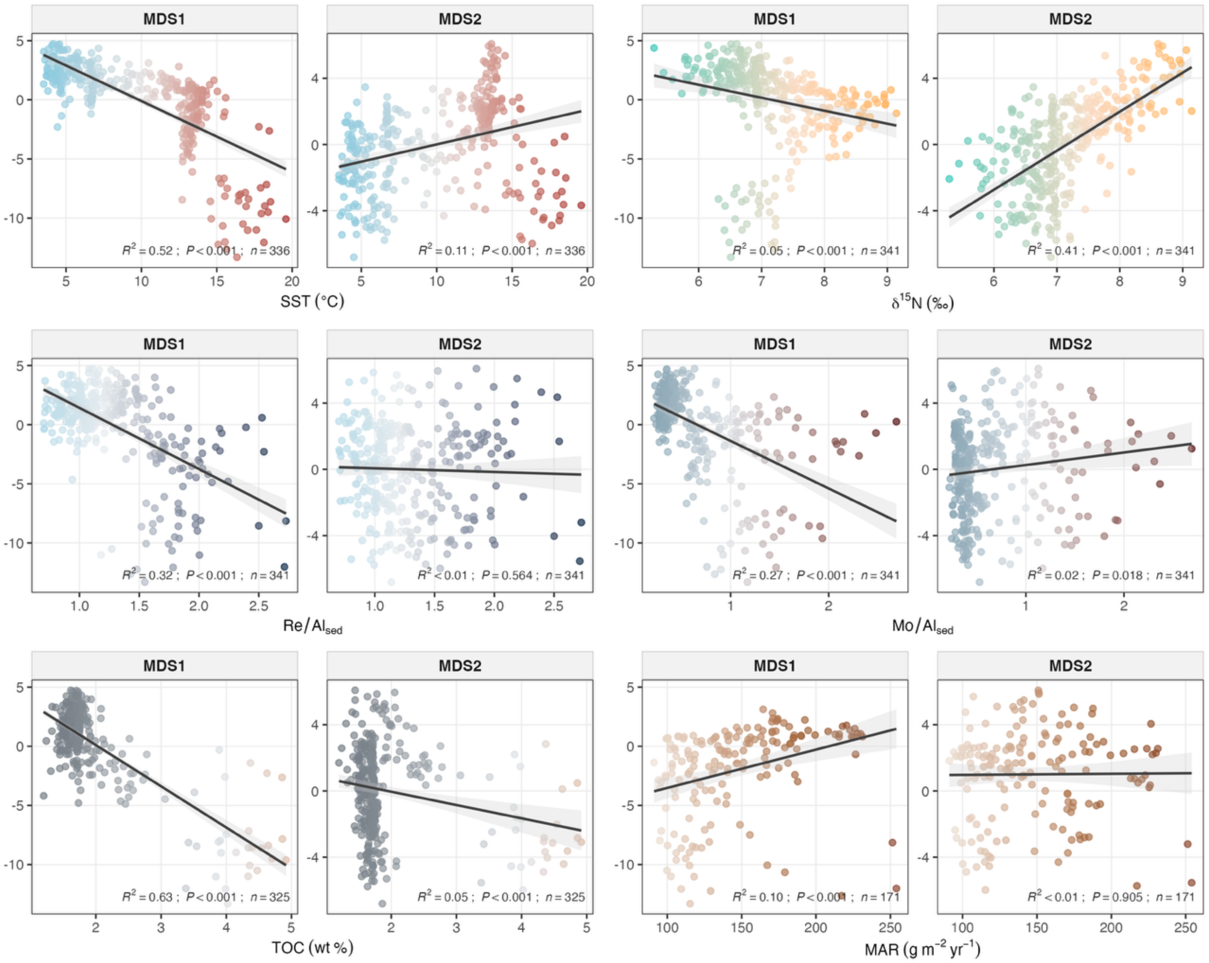


**Fig. S7. Relationships between environmental proxy variables and NMDS ordination axes 1 and 2.**

Each panel pair shows linear relationships between MDS1 (left) and MDS2 (right), for sea surface temperature (SST, ˚C), $\delta^{15}N$ ( ‰), Re/Al, Mo/Al, TOC (wt %), and sedimentary MAR (g $m^{-2}$ $yr^{-1}$). Points are colored by the value of each proxy variable; lines show linear fits with 95% shaded confidence intervals; $R^2$ and p values as well as sample size (n) is shown for each, representing a single-variable linear model for each MDS axis and proxy variable.

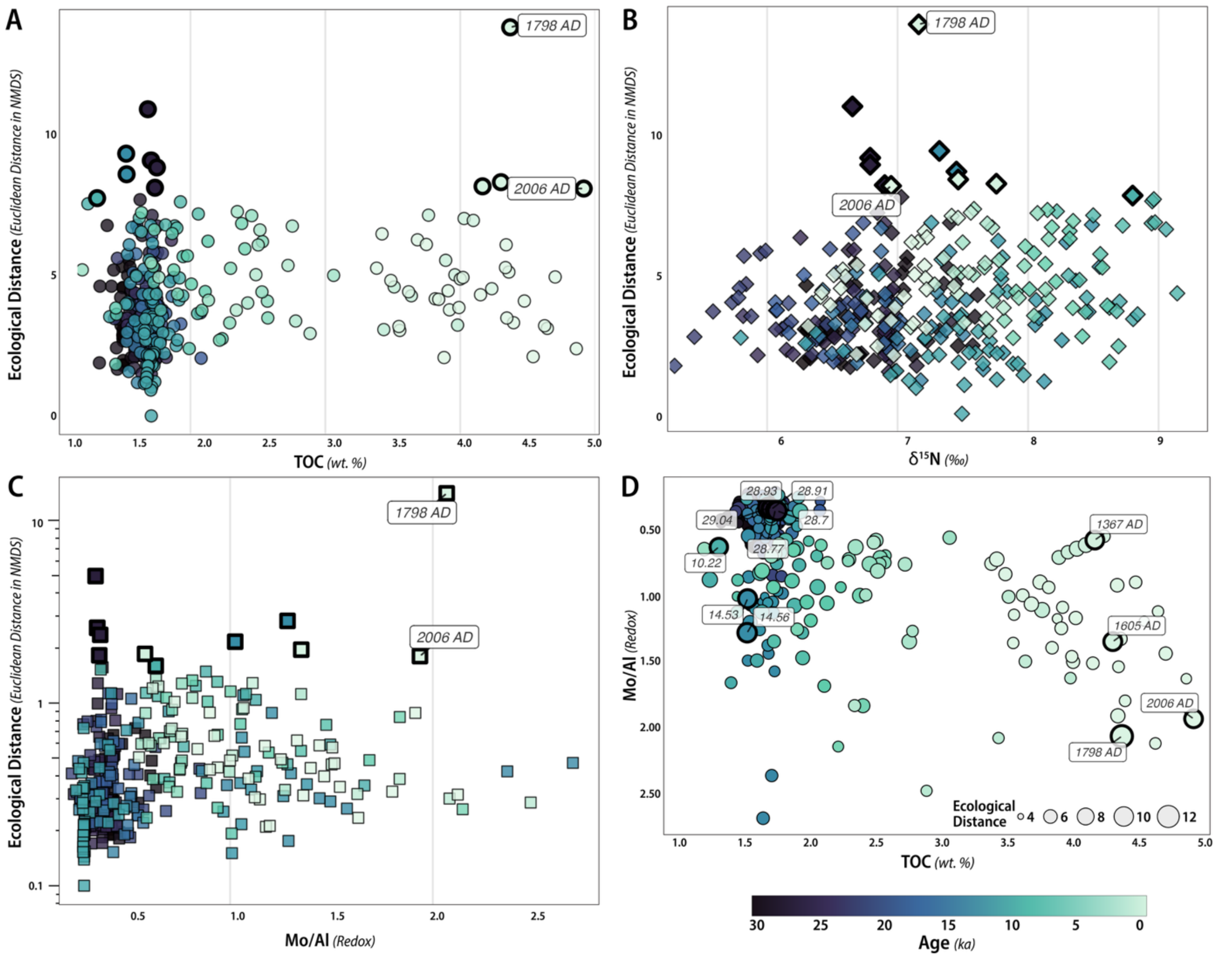


**Fig. S8. Ecological distance, for TOC, δ$^{15}$N, and Mo/Al through time and relationship between TOC and Mo/Al, and ecological distance.**

Ecological distance (Euclidean dissimilarity in NMDS) plotted against values of (**A**) TOC, (**B**) δ$^{15}$N, and (**C**) Mo/Al (redox proxy); (**D**) relationship between TOC and Mo/Al, with point size corresponding to values for ecological distance. Points are colored by age (ka), and heavy-bordered points represent ecological distance outliers, labeled with age (ka) or calendar year (AD). In panel D, the relationship between TOC, Mo/Al, and ecological distance suggests that the highest values of both of the past 34 kya emerged during the late Holocene and are associated with heightened variability.

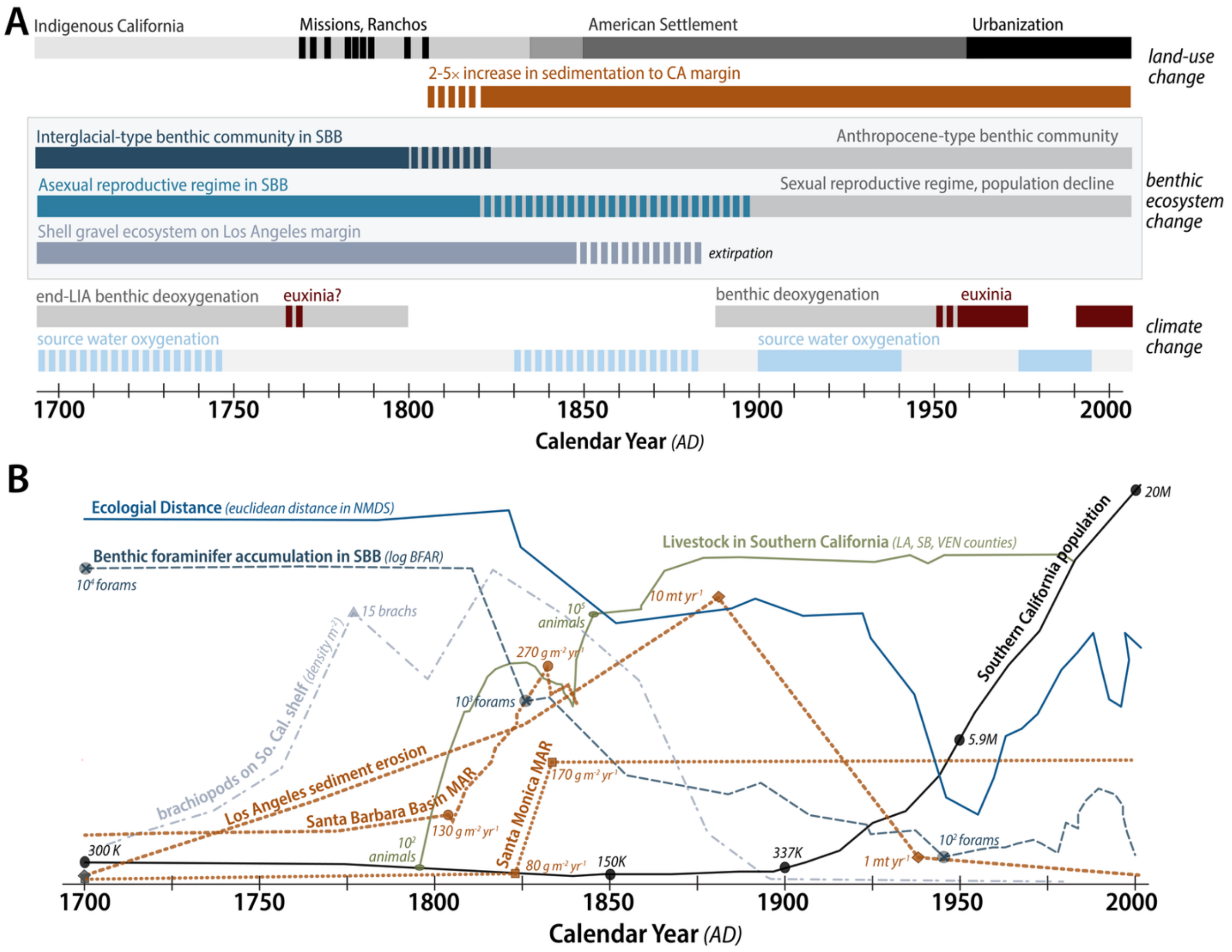


**Fig. S9. Timeline of land-use, climate, and benthic marine ecosystem changes in Southern California, 1700 AD to present.**

**(A)** Categorial timeline of land-use, benthic ecosystem, and climate changes; hatched bars indicate transitional intervals. **(B)** Time series of key ecological and anthropogenic indicators from 1700 to 2000 AD. Ecological distance (Euclidean distance in NMDS space; blue) tracks assemblage dissimilarity from a pre-disturbance baseline. Benthic foraminifer accumulation rate in SBB (log BFAR; dark blue dashed line) and brachiopod density on the Southern California shelf (gray dash-dot) reflect changing benthic community structure. Mass accumulation rates (MAR, g $m^{-2}$ $yr^{-1}$) are shown for Santa Barbara Basin (brown, dotted and dashed) and the Santa Monica Basin (brown, dotted), along with estimated rates of Los Angeles sediment erosion (brown, dashed; in kt $yr^{-1}$), all increasing markedly after ~1800 AD. Livestock numbers in Los Angeles, Santa Barbara, and Ventura counties (olive green) and Southern California human population (black) are overlain to illustrate the coupling between land-use intensification and marine ecological changes. Annotated values provide reference magnitudes at key time points.

**Table S1.**
Data sources for benthic community composition and environmental variables used in this study.

| Variable | Proxy | Ref(s) | Core(s) | Age Range |
|---|---|---|---|---|
| Benthic community composition | Foraminiferal accumulation rate (no. foraminifera x 100 cm-2 yr -1) | Hill et al. 2006, Ohkushi et al. 2013, Moffitt et al. 2014 | MD02-2503 | 34 ka - 0.1 ka |
| | | Kahanamoku et al. 2024 | MV1012-BC1 | 1834 AD - 2008 AD |
| Sea Surface Temperature (SST) | $\delta^{18}O$ | White et al. 2012 | ODP 893, MV0508-11JPC, MV0508-16JPC, MV0508-20JPC | 59.884 ka - 0.358 ka |
| | | | | (57,934 BC to 1592 AD) |

| | | | | |
|---|---|---|---|---|
| | Uk37 (alkenones) | Zhao et al. 2000; NOAA NCEI source url | SABA87-1, SABA87-2, SABA88-1 | 0.652 ka - 0.009 ka |
| | | | | (1298 BC - 1941 AD |
| | Direct measurement | CalCOFI | N/A; bottle database (https://calcofi.org/data/oceanographic-data/bottle-database/) | 1953 AD - present |
| Organic carbon deposition (TOC) | TOC (wt %) | Ivanochko and Pedersen 2004 | ODP 893 | 48.9 ka to 0.357 ka |
| | | Wang et al. 2019 | SPR0901-03KC | 2.1 ka - 1910 AD |

| | | | | |
|---|---|---|---|---|
| | | Xu et al. 2024 | SBB-190629 MUC | 1934 AD - 2018 AD |
| Denitrification | δ$^{15}$N | Emmer and Thunnel 2000 | ODP 893 | 48.6 ka - 1927 AD |
| | | Xu et al. 2024 | SBB-190629 MUC | 1934 AD - 2018 AD |
| Redox | Mo/Al | Wang et al. 2017 | SPR0901-04BC | 1759 AD - 2008 AD |
| | | Ivanochko and Pedersen 2004 | ODP 893 | 48.5 ka to 1807 AD |

**Table S2.**
Sediment cores, sample ages, cluster group, values on MDS axes 1-4, and euclidean dissimilarities.

| Core | Age (ya) | Calendar Age (AD) | Cluster | MDS 1 | MDS 2 | MDS 3 | MDS 4 | sample age | previous sample age | Euclidean dissimilarity |
|---|---|---|---|---|---|---|---|---|---|---|
| MV1012-BC1 | -57 | 2007 | Anthropocene | -2.282 | 0.114 | -1.293 | -1.671 | -57 | -58 | 3.552 |
| MV1012-BC1 | -56 | 2006 | Anthropocene | -9.617 | -3.089 | -0.228 | -3.211 | -56 | -57 | 8.955 |
| MV1012-BC1 | -54 | 2004 | Anthropocene | -7.845 | -2.841 | -0.785 | -2.287 | -54 | -56 | 2.647 |
| MV1012-BC1 | -53 | 2003 | Anthropocene | -8.97 | -4.278 | 1.147 | -2.797 | -53 | -54 | 4.771 |
| MV1012-BC1 | -52 | 2002 | Anthropocene | -11.023 | -4.549 | 0.374 | -1.077 | -52 | -53 | 4.591 |
| MV1012-BC1 | -51 | 2001 | Anthropocene | -13.307 | -5.988 | -0.042 | -1.556 | -51 | -52 | 5.451 |
| MV1012-BC1 | -48 | 1998 | Anthropocene | -10.996 | -4.884 | 0.689 | -2.241 | -48 | -51 | 5.841 |
| MV1012-BC1 | -46 | 1996 | Anthropocene | -10.502 | -3.141 | -0.335 | -1.316 | -46 | -48 | 4.525 |
| MV1012-BC1 | -44 | 1994 | Anthropocene | -8.624 | -2.022 | -0.361 | -0.763 | -44 | -46 | 2.334 |
| MV1012-BC1 | -40 | 1990 | Anthropocene | -12.065 | -4.598 | -0.07 | -1.324 | -40 | -44 | 4.267 |

| | | | | | | | | | | |
|---|---|---|---|---|---|---|---|---|---|---|
| MV1012-BC1 | -39 | 1989 | Anthropocene | -7.444 | -1.697 | -1.591 | -0.584 | -39 | -40 | 5.392 |
| MV1012-BC1 | -37 | 1987 | Anthropocene | -7.974 | -2.367 | -0.237 | 0.491 | -37 | -39 | 4.929 |
| MV1012-BC1 | -35 | 1985 | Anthropocene | -8.386 | -4.417 | -0.004 | 0.252 | -35 | -37 | 3.448 |
| MV1012-BC1 | -30 | 1980 | Anthropocene | -10.084 | -3.679 | -0.936 | -0.474 | -30 | -35 | 5.665 |
| MV1012-BC1 | -28 | 1978 | Anthropocene | -7.168 | -2.846 | -0.882 | -0.832 | -28 | -30 | 3.855 |
| MV1012-BC1 | -24 | 1974 | Anthropocene | -9.494 | -3.631 | 0.639 | -1.402 | -24 | -28 | 5.472 |
| MV1012-BC1 | -18 | 1968 | Anthropocene | -8.594 | -3.052 | -1.713 | -1.41 | -18 | -24 | 6.739 |
| MV1012-BC1 | -16 | 1966 | Anthropocene | -8.552 | -4.035 | -1.68 | -0.448 | -16 | -18 | 3.67 |
| MV1012-BC1 | -13 | 1963 | Anthropocene | -8.455 | -2.485 | -0.983 | -1.275 | -13 | -16 | 3.522 |
| MV1012-BC1 | -3 | 1953 | Anthropocene | -9.012 | -1.61 | -0.563 | -0.044 | -3 | -13 | 4.178 |
| MV1012-BC1 | 6 | 1944 | Anthropocene | -8.093 | -0.997 | -1.135 | -0.792 | 6 | -3 | 2.298 |
| MV1012-BC1 | 18 | 1932 | Anthropocene | -7.068 | 0.42 | -1.187 | 0.406 | 18 | 6 | 4.612 |

| | | | | | | | | | | |
|---|---|---|---|---|---|---|---|---|---|---|
| MV1012-BC1 | 21 | 1929 | Anthropocene | -12.236 | -2.984 | 0.34 | -0.192 | 21 | 18 | 6.751 |
| MV1012-BC1 | 36 | 1914 | Anthropocene | -6.642 | -1.398 | -0.809 | -0.704 | 36 | 21 | 6.935 |
| MV1012-BC1 | 50 | 1900 | Anthropocene | -10.061 | -2.335 | -1.354 | -0.367 | 50 | 36 | 4.888 |
| MV1012-BC1 | 59 | 1891 | Anthropocene | -10.661 | -2.937 | 0.06 | -0.402 | 59 | 50 | 3.366 |
| MV1012-BC1 | 88 | 1862 | Anthropocene | -12.632 | -5.74 | 1.158 | 1.339 | 88 | 59 | 5.333 |
| MD02-2503 | 114 | 1836 | Anthropocene | -8.142 | -3.221 | -0.513 | 1.469 | 114 | 88 | 7.168 |
| MD02-2503 | 116 | 1834 | Anthropocene | -12.02 | -5.559 | 0.714 | 1.782 | 116 | 114 | 5.818 |
| MD02-2503 | 152 | 1798 | Anthropocene | -1.408 | 2.831 | -2.515 | 2.175 | 152 | 116 | 15.303 |
| MD02-2503 | 345 | 1605 | interglacial | -7.485 | -0.013 | -0.586 | 0.835 | 345 | 152 | 9.205 |
| MD02-2503 | 442 | 1508 | interglacial | -6.644 | 1.272 | 0.825 | 3.212 | 442 | 345 | 5.009 |
| MD02-2503 | 583 | 1367 | interglacial | -1.045 | 2.16 | -2.315 | -0.014 | 583 | 442 | 9.041 |
| MD02-2503 | 761 | 1189 | interglacial | -6.097 | 0.847 | 0.332 | 0.355 | 761 | 583 | 7.698 |
| MD02-2503 | 894 | 1056 | interglacial | -0.851 | 1.4 | -1.463 | 0.163 | 894 | 761 | 7.774 |
| MD02-2503 | 1020 | 930 | interglacial | -2.974 | 2.127 | -3.72 | 1.117 | 1020 | 894 | 5.557 |

| | | | | | | | | | | |
|---|---|---|---|---|---|---|---|---|---|---|
| MD02-2503 | 1206 | 744 | interglacial | -0.309 | 2.203 | -2.194 | -0.356 | 1206 | 1020 | 6.128 |
| MD02-2503 | 1354 | 596 | interglacial | -5.217 | 2.038 | -1.792 | 2.505 | 1354 | 1206 | 7.905 |
| MD02-2503 | 1488 | 462 | interglacial | -2.837 | 2.335 | -1.848 | 2.768 | 1488 | 1354 | 4.73 |
| MD02-2503 | 1651 | 299 | interglacial | -3.897 | 0.84 | -0.728 | 0.174 | 1651 | 1488 | 5.079 |
| MD02-2503 | 1822 | 128 | interglacial | -2.588 | 1.47 | -1.677 | 0.168 | 1822 | 1651 | 3.577 |
| MD02-2503 | 1948 | 2 | interglacial | -1.646 | 1.685 | -1.07 | 0.187 | 1948 | 1822 | 3.399 |
| MD02-2503 | 2059 | -109 | interglacial | -0.701 | 2.018 | -2.051 | 0.158 | 2059 | 1948 | 3.245 |
| MD02-2503 | 2223 | -273 | interglacial | -2.905 | 1.561 | -1.85 | -0.321 | 2223 | 2059 | 4.165 |
| MD02-2503 | 2408 | -458 | interglacial | -3.5 | 2.088 | -0.91 | 1.99 | 2408 | 2223 | 5.562 |
| MD02-2503 | 2557 | -607 | interglacial | -2.855 | 0.761 | -1.159 | -0.923 | 2557 | 2408 | 5.927 |
| MD02-2503 | 2661 | -711 | interglacial | -0.691 | 1.443 | -1.195 | -1.051 | 2661 | 2557 | 3.736 |
| MD02-2503 | 2883 | -933 | interglacial | -2.365 | 2.714 | -0.723 | 3.372 | 2883 | 2661 | 7.476 |
| MD02-2503 | 3002 | -1052 | interglacial | 0.639 | 2.166 | -1.881 | 0.227 | 3002 | 2883 | 7.323 |
| MD02-2503 | 3113 | -1163 | interglacial | -1.701 | 1.571 | -2.247 | 0.907 | 3113 | 3002 | 4.448 |
| MD02-2503 | 3321 | -1371 | interglacial | -3.665 | 3.783 | 1.616 | 4.635 | 3321 | 3113 | 8.045 |
| MD02-2503 | 3477 | -1527 | interglacial | -4.301 | 1.676 | -1.921 | 0.946 | 3477 | 3321 | 6.732 |
| MD02-2503 | 3625 | -1675 | interglacial | -4.455 | 0.937 | -1.043 | 2.304 | 3625 | 3477 | 5.535 |

| | | | | | | | | | | |
|---|---|---|---|---|---|---|---|---|---|---|
| MD02-2503 | 3766 | -1816 | interglacial | -3.723 | 1.284 | 0.485 | -0.425 | 3766 | 3625 | 5.459 |
| MD02-2503 | 3885 | -1935 | interglacial | -4.188 | 1.694 | -1.059 | 1.553 | 3885 | 3766 | 5.751 |
| MD02-2503 | 4048 | -2098 | interglacial | -1.887 | 0.944 | -1.033 | 4.558 | 4048 | 3885 | 6.031 |
| MD02-2503 | 4197 | -2247 | interglacial | -2.21 | 2.895 | -1.724 | 2.753 | 4197 | 4048 | 4.556 |
| MD02-2503 | 4345 | -2395 | interglacial | -4.877 | 0.836 | 0.385 | 2.121 | 4345 | 4197 | 7.885 |
| MD02-2503 | 4471 | -2521 | interglacial | -3.188 | 0.347 | -0.718 | 2.663 | 4471 | 4345 | 4.883 |
| MD02-2503 | 4650 | -2700 | interglacial | -3.722 | 1.345 | -1.483 | 2.448 | 4650 | 4471 | 4.52 |
| MD02-2503 | 4798 | -2848 | interglacial | -1.932 | 1.395 | -1.387 | 1.964 | 4798 | 4650 | 3.664 |
| MD02-2503 | 4946 | -2996 | interglacial | -1.275 | 0.987 | -1.318 | 4.849 | 4946 | 4798 | 5.813 |
| MD02-2503 | 5095 | -3145 | interglacial | -2.301 | 1.484 | -0.074 | 0.422 | 5095 | 4946 | 6.862 |
| MD02-2503 | 5280 | -3330 | interglacial | -2.29 | 1.689 | 0.475 | -0.031 | 5280 | 5095 | 3.858 |
| MD02-2503 | 5421 | -3471 | interglacial | -2.425 | 1.99 | -1.812 | 0.362 | 5421 | 5280 | 4.799 |
| MD02-2503 | 5548 | -3598 | interglacial | -1.446 | 2.131 | -1.42 | 1.576 | 5548 | 5421 | 2.982 |
| MD02-2503 | 5681 | -3731 | interglacial | -0.642 | 1.109 | -1.643 | 4.07 | 5681 | 5548 | 6.579 |
| MD02-2503 | 5815 | -3865 | interglacial | -4.643 | 0.988 | -0.765 | 0.396 | 5815 | 5681 | 7.577 |
| MD02-2503 | 5963 | -4013 | interglacial | -3.276 | 1.888 | -0.67 | 0.684 | 5963 | 5815 | 4.159 |
| MD02-2503 | 6089 | -4139 | interglacial | -4.804 | -1.66 | 1.532 | 3.827 | 6089 | 5963 | 7.331 |

| | | | | | | | | | | |
|---|---|---|---|---|---|---|---|---|---|---|
| MD02-2503 | 6230 | -4280 | interglacial | -2.484 | 2.108 | 0.793 | -0.407 | 6230 | 6089 | 8.007 |
| MD02-2503 | 6423 | -4473 | interglacial | -1.368 | 1.821 | -0.909 | 1.742 | 6423 | 6230 | 5.748 |
| MD02-2503 | 6572 | -4622 | interglacial | -1.722 | 0.755 | 1.276 | 2.073 | 6572 | 6423 | 5.889 |
| MD02-2503 | 6705 | -4755 | interglacial | -0.348 | 3.92 | 2.094 | 1.561 | 6705 | 6572 | 5.192 |
| MD02-2503 | 6876 | -4926 | interglacial | 0.58 | 4.357 | 0.939 | -0.983 | 6876 | 6705 | 4.897 |
| MD02-2503 | 7047 | -5097 | interglacial | -0.976 | 3.975 | 0.149 | 2.111 | 7047 | 6876 | 6.308 |
| MD02-2503 | 7173 | -5223 | interglacial | -0.114 | 3.935 | 0.911 | 2.222 | 7173 | 7047 | 3.041 |
| MD02-2503 | 7321 | -5371 | interglacial | 0.187 | 2.382 | -2.054 | -0.167 | 7321 | 7173 | 6.88 |
| MD02-2503 | 7529 | -5579 | interglacial | -0.236 | 4.671 | 1.119 | 1.031 | 7529 | 7321 | 7.182 |
| MD02-2503 | 7648 | -5698 | interglacial | -0.731 | 5.498 | 0.291 | 1.299 | 7648 | 7529 | 2.036 |
| MD02-2503 | 7767 | -5817 | interglacial | -1.897 | 6.065 | 0.75 | -0.782 | 7767 | 7648 | 4.251 |
| MD02-2503 | 7908 | -5958 | interglacial | -0.437 | 4.654 | 1.218 | 0.811 | 7908 | 7767 | 4.002 |
| MD02-2503 | 8056 | -6106 | interglacial | 0.444 | 4.055 | 1.179 | 2.256 | 8056 | 7908 | 2.653 |
| MD02-2503 | 8323 | -6373 | interglacial | -2.993 | 4.982 | 1.966 | 0.645 | 8323 | 8056 | 5.259 |
| MD02-2503 | 8501 | -6551 | interglacial | -2.411 | 3.258 | 0.931 | 1.332 | 8501 | 8323 | 6.274 |
| MD02-2503 | 8650 | -6700 | interglacial | -1.844 | 5.009 | 1.165 | -2.344 | 8650 | 8501 | 7.954 |
| MD02-2503 | 8806 | -6856 | interglacial | -0.38 | 5.941 | 0.576 | -0.476 | 8806 | 8650 | 4.031 |

| | | | | | | | | | | |
|---|---|---|---|---|---|---|---|---|---|---|
| MD02 -2503 | 8932 | -6982 | intergl acial | -1.376 | 5.827 | 1.351 | -1.921 | 8932 | 8806 | 4.167 |
| MD02 -2503 | 9095 | -7145 | intergl acial | -1.449 | 5.272 | 0.535 | 2.406 | 9095 | 8932 | 7.397 |
| MD02 -2503 | 9258 | -7308 | intergl acial | -0.442 | 4.265 | 1.865 | -2.078 | 9258 | 9095 | 8.43 |
| MD02 -2503 | 9392 | -7442 | intergl acial | -0.334 | 4.878 | -0.304 | -3.144 | 9392 | 9258 | 7.522 |
| MD02 -2503 | 9540 | -7590 | intergl acial | 0.423 | 2.463 | -2.068 | -0.12 | 9540 | 9392 | 7.796 |
| MD02 -2503 | 9689 | -7739 | intergl acial | 0.161 | 5.087 | 1.339 | -1.427 | 9689 | 9540 | 7.482 |
| MD02 -2503 | 9815 | -7865 | intergl acial | -1.151 | 2.02 | 0.512 | -0.945 | 9815 | 9689 | 4.733 |
| MD02 -2503 | 9986 | -8036 | intergl acial | 0.798 | 2.638 | -2.169 | -0.436 | 9986 | 9815 | 5.432 |
| MD02 -2503 | 10223 | -8273 | intergl acial | -2.108 | 4.055 | 2.443 | -2.058 | 10223 | 9986 | 8.576 |
| MD02 -2503 | 10372 | -8422 | intergl acial | 0.451 | 2.583 | -1.709 | -0.154 | 10372 | 10223 | 8.352 |
| MD02 -2503 | 10520 | -8570 | intergl acial | 0.823 | 5.689 | 1.106 | -1.521 | 10520 | 10372 | 7.247 |
| MD02 -2503 | 10668 | -8718 | intergl acial | -0.275 | 4.753 | 1.139 | -0.732 | 10668 | 10520 | 3.908 |
| MD02 -2503 | 10787 | -8837 | intergl acial | 0.664 | 2.414 | -0.975 | 0.08 | 10787 | 10668 | 5.505 |
| MD02 -2503 | 10965 | -9015 | intergl acial | 1.27 | 5.361 | 0.193 | -0.703 | 10965 | 10787 | 4.796 |
| MD02 -2503 | 11114 | -9164 | intergl acial | 0.029 | 5.071 | 1.438 | -1.765 | 11114 | 10965 | 3.722 |
| MD02 -2503 | 11255 | -9305 | intergl acial | 0.839 | 4.897 | -1.161 | -0.567 | 11255 | 11114 | 4.563 |
| MD02 -2503 | 11411 | -9461 | intergl acial | -0.396 | 2.021 | -2.163 | -1.096 | 11411 | 11255 | 4.407 |

| | | | | | | | | | | |
|---|---|---|---|---|---|---|---|---|---|---|
| MD02-2503 | 11514 | -9564 | interglacial | -0.138 | 2.591 | -1.767 | -0.636 | 11514 | 11411 | 3.233 |
| MD02-2503 | 11522 | -9572 | interglacial | -1.465 | 2.785 | -2.849 | 0.707 | 11522 | 11514 | 3.364 |
| MD02-2503 | 11529 | -9579 | interglacial | 0.173 | 2.144 | -1.893 | 0.551 | 11529 | 11522 | 3.816 |
| MD02-2503 | 11537 | -9587 | interglacial | 0.711 | 2.592 | -2.305 | -0.53 | 11537 | 11529 | 2.983 |
| MD02-2503 | 11544 | -9594 | interglacial | 0.642 | 2.581 | -2.291 | -0.593 | 11544 | 11537 | 1.313 |
| MD02-2503 | 11552 | -9602 | interglacial | 0.834 | 2.595 | -2.251 | -0.441 | 11552 | 11544 | 1.523 |
| MD02-2503 | 11559 | -9609 | interglacial | 0.397 | 2.241 | -1.217 | -0.406 | 11559 | 11552 | 1.684 |
| MD02-2503 | 11566 | -9616 | interglacial | 0.74 | 2.675 | -1.861 | -0.506 | 11566 | 11559 | 1.523 |
| MD02-2503 | 11574 | -9624 | interglacial | 0.762 | 2.088 | -1.994 | 0.24 | 11574 | 11566 | 1.967 |
| MD02-2503 | 11581 | -9631 | interglacial | 0.575 | 1.74 | -1.12 | -0.323 | 11581 | 11574 | 1.967 |
| MD02-2503 | 11589 | -9639 | glacial | 1.288 | -1.278 | -0.793 | 0.665 | 11589 | 11581 | 4.794 |
| MD02-2503 | 11596 | -9646 | interglacial | 0.927 | 2.35 | -2.205 | -0.459 | 11596 | 11589 | 6.147 |
| MD02-2503 | 11604 | -9654 | interglacial | 0.946 | 2.153 | -1.173 | -0.3 | 11604 | 11596 | 2.373 |
| MD02-2503 | 11611 | -9661 | interglacial | 1.255 | 2.266 | -2.039 | 0.221 | 11611 | 11604 | 2.32 |
| MD02-2503 | 11618 | -9668 | interglacial | 0.782 | 2.523 | -2.233 | -0.33 | 11618 | 11611 | 2.246 |
| MD02-2503 | 11626 | -9676 | interglacial | 0.809 | 2.558 | -2.212 | -0.432 | 11626 | 11618 | 1.017 |
| MD02-2503 | 11633 | -9683 | interglacial | 0.809 | 2.558 | -2.212 | -0.432 | 11633 | 11626 | 0 |

| | | | | | | | | | | |
|---|---|---|---|---|---|---|---|---|---|---|
| MD02 -2503 | 11641 | -9691 | intergl acial | 1.047 | 2.29 | -2.037 | -0.067 | 11641 | 11633 | 1.133 |
| MD02 -2503 | 11648 | -9698 | intergl acial | 1.097 | 2.196 | -2.332 | 0.335 | 11648 | 11641 | 2.015 |
| MD02 -2503 | 11653 | -9703 | glacial | 1.495 | 0.347 | -1.424 | 1.143 | 11653 | 11648 | 3.797 |
| MD02 -2503 | 11655 | -9705 | intergl acial | 0.806 | 0.215 | -0.798 | 2.292 | 11655 | 11653 | 3.892 |
| MD02 -2503 | 11663 | -9713 | intergl acial | 1.377 | 0.955 | -0.844 | 1.55 | 11663 | 11655 | 2.412 |
| MD02 -2503 | 11670 | -9720 | intergl acial | 1.765 | 0.693 | -1.477 | 1.225 | 11670 | 11663 | 2.402 |
| MD02 -2503 | 11678 | -9728 | glacial | 1.678 | 0.494 | -0.798 | 1.435 | 11678 | 11670 | 1.979 |
| MD02 -2503 | 11685 | -9735 | intergl acial | 1.014 | 0.734 | -1.39 | 0.522 | 11685 | 11678 | 2.494 |
| MD02 -2503 | 11693 | -9743 | intergl acial | 1 | 0.755 | -1.439 | 0.427 | 11693 | 11685 | 1.344 |
| MD02 -2503 | 11781 | -9831 | glacial | 0.683 | -0.775 | -1.141 | -0.113 | 11781 | 11693 | 2.859 |
| MD02 -2503 | 11824 | -9874 | glacial | 2.229 | -0.982 | -1.575 | 1.41 | 11824 | 11781 | 3.28 |
| MD02 -2503 | 11930 | -9980 | glacial | 2.114 | -2.958 | -0.104 | -1.262 | 11930 | 11824 | 5.289 |
| MD02 -2503 | 12037 | - 10087 | glacial | 1.978 | -2.67 | -0.863 | 0.345 | 12037 | 11930 | 2.992 |
| MD02 -2503 | 12138 | - 10188 | glacial | 1.271 | -3.645 | -0.884 | -0.038 | 12138 | 12037 | 2.766 |
| MD02 -2503 | 12240 | - 10290 | glacial | 3.118 | -4.297 | -0.721 | -0.203 | 12240 | 12138 | 3.221 |
| MD02 -2503 | 12346 | - 10396 | glacial | 2.814 | -3.741 | 0.09 | 0.208 | 12346 | 12240 | 2.463 |
| MD02 -2503 | 12517 | - 10567 | glacial | 1.655 | -2.81 | 0.708 | -0.899 | 12517 | 12346 | 3.552 |

| | | | | | | | | | | |
|---|---|---|---|---|---|---|---|---|---|---|
| MD02-2503 | 12672 | -10722 | glacial | 0.727 | -2.792 | 3.426 | 1.648 | 12672 | 12517 | 4.599 |
| MD02-2503 | 12800 | -10850 | glacial | 2.231 | -0.794 | 1.979 | 4.011 | 12800 | 12672 | 5.495 |
| MD02-2503 | 12846 | -10896 | interglacial | 1.337 | 2.976 | 0.07 | 0.1 | 12846 | 12800 | 7.506 |
| MD02-2503 | 12934 | -10984 | interglacial | -1.632 | 1.16 | 3.389 | 0.249 | 12934 | 12846 | 6.062 |
| MD02-2503 | 13055 | -11105 | interglacial | 0.754 | 3.335 | -0.445 | 0.192 | 13055 | 12934 | 6.799 |
| MD02-2503 | 13175 | -11225 | interglacial | 1.324 | 3.167 | 0.464 | 0.271 | 13175 | 13055 | 2.397 |
| MD02-2503 | 13285 | -11335 | interglacial | 0.597 | 2.692 | -1.57 | -0.285 | 13285 | 13175 | 4.467 |
| MD02-2503 | 13367 | -11417 | interglacial | 0.472 | 3.2 | -0.055 | -0.393 | 13367 | 13285 | 2.892 |
| MD02-2503 | 13494 | -11544 | interglacial | 0.245 | 2.83 | -0.142 | -0.365 | 13494 | 13367 | 1.269 |
| MD02-2503 | 13603 | -11653 | interglacial | -0.67 | 2.643 | 1.485 | -0.983 | 13603 | 13494 | 3.753 |
| MD02-2503 | 13730 | -11780 | interglacial | 0.232 | 3.476 | 0.274 | -0.755 | 13730 | 13603 | 3.493 |
| MD02-2503 | 13823 | -11873 | interglacial | 0.241 | 1.833 | -0.715 | -0.984 | 13823 | 13730 | 3.956 |
| MD02-2503 | 13933 | -11983 | interglacial | -0.449 | 2.307 | 0.899 | -0.093 | 13933 | 13823 | 4.006 |
| MD02-2503 | 14042 | -12092 | interglacial | -1.287 | 3.428 | 1.506 | 2.853 | 14042 | 13933 | 5.777 |
| MD02-2503 | 14158 | -12208 | interglacial | 0.304 | 2.208 | -0.308 | -0.542 | 14158 | 14042 | 7.424 |
| MD02-2503 | 14240 | -12290 | interglacial | -0.686 | 4.055 | 2.258 | -0.687 | 14240 | 14158 | 5.227 |
| MD02-2503 | 14317 | -12367 | interglacial | 0.858 | 2.817 | -1.678 | -0.483 | 14317 | 14240 | 6.741 |

| | | | | | | | | | | |
|---|---|---|---|---|---|---|---|---|---|---|
| MD02 -2503 | 14355 | - 12405 | intergl acial | 0.81 | 2.976 | -1.298 | -0.726 | 14355 | 14317 | 1.747 |
| MD02 -2503 | 14405 | - 12455 | intergl acial | 0.94 | 0.933 | -0.009 | 0.263 | 14405 | 14355 | 4.384 |
| MD02 -2503 | 14443 | - 12493 | intergl acial | 0.809 | 2.558 | -2.212 | -0.432 | 14443 | 14405 | 4.476 |
| MD02 -2503 | 14470 | - 12520 | intergl acial | -1.719 | 0.084 | -0.891 | -2.116 | 14470 | 14443 | 6.359 |
| MD02 -2503 | 14531 | - 12581 | glacial | 0.902 | -3.929 | 1.276 | 5.107 | 14531 | 14470 | 9.517 |
| MD02 -2503 | 14564 | - 12614 | glacial | 1.595 | 0.382 | -2.535 | -0.459 | 14564 | 14531 | 10.328 |
| MD02 -2503 | 14591 | - 12641 | glacial | 1.164 | -0.15 | -1.974 | 0.419 | 14591 | 14564 | 3.287 |
| MD02 -2503 | 14712 | - 12762 | glacial | 1.262 | -0.18 | -1.064 | -0.214 | 14712 | 14591 | 3.007 |
| MD02 -2503 | 14860 | - 12910 | glacial | 2.339 | -0.659 | -2.285 | 0.375 | 14860 | 14712 | 3.41 |
| MD02 -2503 | 14965 | - 13015 | intergl acial | 0.251 | 1.249 | -2.725 | -0.778 | 14965 | 14860 | 4.791 |
| MD02 -2503 | 15078 | - 13128 | glacial | 0.913 | -0.886 | -2.259 | 0.649 | 15078 | 14965 | 4.458 |
| MD02 -2503 | 15218 | - 13268 | glacial | 2.33 | -0.71 | -2.353 | 0.541 | 15218 | 15078 | 3.922 |
| MD02 -2503 | 15359 | - 13409 | glacial | 1.582 | -2.154 | -2.074 | -0.062 | 15359 | 15218 | 2.967 |
| MD02 -2503 | 15527 | - 13577 | glacial | 1.562 | -0.775 | -3.026 | 0.287 | 15527 | 15359 | 2.49 |
| MD02 -2503 | 15633 | - 13683 | glacial | 2.074 | -1.11 | -2.628 | 0.545 | 15633 | 15527 | 1.311 |
| MD02 -2503 | 15773 | - 13823 | glacial | 1.529 | 0.416 | -2.731 | 0.829 | 15773 | 15633 | 3.116 |
| MD02 -2503 | 15942 | - 13992 | glacial | 1.845 | -0.45 | -3.553 | 0.307 | 15942 | 15773 | 2.936 |

| | | | | | | | | | | |
|---|---|---|---|---|---|---|---|---|---|---|
| MD02 -2503 | 16090 | - 14140 | glacial | 2.011 | -0.129 | -2.3 | -0.98 | 16090 | 15942 | 5.544 |
| MD02 -2503 | 16329 | - 14379 | glacial | 3.006 | -1.403 | -2.616 | -0.057 | 16329 | 16090 | 3.996 |
| MD02 -2503 | 16441 | - 14491 | glacial | 1.728 | 0.27 | -2.96 | 0.092 | 16441 | 16329 | 3.53 |
| MD02 -2503 | 16553 | - 14603 | glacial | 1.582 | -2.193 | -2.107 | 0.85 | 16553 | 16441 | 4.861 |
| MD02 -2503 | 16694 | - 14744 | glacial | 2.664 | -1.94 | -3.909 | 0.891 | 16694 | 16553 | 3.058 |
| MD02 -2503 | 16849 | - 14899 | glacial | 1.331 | -1.866 | -2.264 | 0.247 | 16849 | 16694 | 3.465 |
| MD02 -2503 | 16954 | - 15004 | glacial | 1.42 | -1.648 | -2.694 | -0.303 | 16954 | 16849 | 2.363 |
| MD02 -2503 | 17102 | - 15152 | glacial | 2.45 | -1.99 | -2.774 | 0.332 | 17102 | 16954 | 1.996 |
| MD02 -2503 | 17235 | - 15285 | glacial | 2.249 | -1.673 | -1.597 | 0.111 | 17235 | 17102 | 2.457 |
| MD02 -2503 | 17376 | - 15426 | glacial | 2.728 | -2.304 | -1.663 | -0.086 | 17376 | 17235 | 2.74 |
| MD02 -2503 | 17516 | - 15566 | glacial | 2.687 | -4.083 | -1.097 | 0.135 | 17516 | 17376 | 3.608 |
| MD02 -2503 | 17657 | - 15707 | glacial | 2.528 | -4.986 | -0.104 | -0.814 | 17657 | 17516 | 2.745 |
| MD02 -2503 | 17776 | - 15826 | glacial | 2.21 | -6.797 | -0.354 | -1.447 | 17776 | 17657 | 4.713 |
| MD02 -2503 | 17889 | - 15939 | glacial | 3.388 | -3.913 | -0.758 | -0.48 | 17889 | 17776 | 7.293 |
| MD02 -2503 | 18149 | - 16199 | glacial | 3.157 | -5.777 | -0.902 | 0.501 | 18149 | 17889 | 4.298 |
| MD02 -2503 | 18275 | - 16325 | glacial | 1.869 | -5.027 | 1.513 | 0.066 | 18275 | 18149 | 6.381 |
| MD02 -2503 | 18367 | - 16417 | glacial | 2.46 | -1.806 | 0.074 | 0.288 | 18367 | 18275 | 5.639 |

| | | | | | | | | | | |
|---|---|---|---|---|---|---|---|---|---|---|
| MD02 -2503 | 18486 | - 16536 | glacial | 2.174 | -5.161 | 0.335 | 0.203 | 18486 | 18367 | 4.439 |
| MD02 -2503 | 18627 | - 16677 | glacial | 2.694 | -4.737 | 1.184 | -1.046 | 18627 | 18486 | 3.506 |
| MD02 -2503 | 18767 | - 16817 | glacial | 1.986 | -3.794 | 0.449 | -1.162 | 18767 | 18627 | 2.782 |
| MD02 -2503 | 18866 | - 16916 | glacial | 2.762 | -2.746 | 0.266 | -0.292 | 18866 | 18767 | 3.428 |
| MD02 -2503 | 18999 | - 17049 | glacial | 2.39 | -4.182 | 0.914 | 0.732 | 18999 | 18866 | 3.657 |
| MD02 -2503 | 19231 | - 17281 | glacial | 3.505 | -4.467 | 0.664 | 0.344 | 19231 | 18999 | 2.434 |
| MD02 -2503 | 19364 | - 17414 | glacial | 2.287 | -2.555 | -0.517 | -0.206 | 19364 | 19231 | 3.79 |
| MD02 -2503 | 19505 | - 17555 | glacial | 1.86 | -4.727 | 0.186 | 0.874 | 19505 | 19364 | 4.346 |
| MD02 -2503 | 19646 | - 17696 | glacial | 1.746 | -4.958 | 1.657 | 0.904 | 19646 | 19505 | 3.17 |
| MD02 -2503 | 19786 | - 17836 | glacial | 0.807 | -5.436 | 0.165 | 2.004 | 19786 | 19646 | 4.071 |
| MD02 -2503 | 19877 | - 17927 | glacial | 0.62 | -3.6 | 1.409 | 3.016 | 19877 | 19786 | 4.379 |
| MD02 -2503 | 20004 | - 18054 | glacial | 2.446 | -4.683 | -0.478 | 1.763 | 20004 | 19877 | 5.45 |
| MD02 -2503 | 20109 | - 18159 | glacial | 2.369 | -4.553 | 1.755 | 0.91 | 20109 | 20004 | 4.059 |
| MD02 -2503 | 20264 | - 18314 | glacial | 3.355 | -3.825 | 1.414 | -0.379 | 20264 | 20109 | 4.037 |
| MD02 -2503 | 20405 | - 18455 | glacial | 3.97 | -5.301 | 2.002 | 0.314 | 20405 | 20264 | 3.625 |
| MD02 -2503 | 20559 | - 18609 | glacial | 3.173 | -4.246 | 2.057 | -0.453 | 20559 | 20405 | 2.272 |
| MD02 -2503 | 20686 | - 18736 | glacial | 2.942 | -3.644 | 2.258 | -0.238 | 20686 | 20559 | 2.05 |

| | | | | | | | | | | |
|---|---|---|---|---|---|---|---|---|---|---|
| MD02 -2503 | 20847 | - 18897 | glacial | 1.584 | -5.125 | -0.295 | 2.359 | 20847 | 20686 | 7.087 |
| MD02 -2503 | 21023 | - 19073 | glacial | 4.355 | -4.512 | -0.805 | 0.682 | 21023 | 20847 | 5.537 |
| MD02 -2503 | 21192 | - 19242 | glacial | 3.24 | -3.178 | 0.422 | 1.709 | 21192 | 21023 | 4.168 |
| MD02 -2503 | 21332 | - 19382 | glacial | 2.101 | -4.562 | 1.292 | -1.467 | 21332 | 21192 | 6.474 |
| MD02 -2503 | 21473 | - 19523 | glacial | 1.865 | -2.092 | -0.875 | 2.312 | 21473 | 21332 | 7.503 |
| MD02 -2503 | 21613 | - 19663 | glacial | 3.058 | -3.166 | 0.381 | 1.262 | 21613 | 21473 | 4.609 |
| MD02 -2503 | 21733 | - 19783 | glacial | 4.227 | -3.738 | 0.195 | 1.568 | 21733 | 21613 | 4.314 |
| MD02 -2503 | 21873 | - 19923 | glacial | 3.991 | -4.333 | 0.473 | 0.69 | 21873 | 21733 | 3.07 |
| MD02 -2503 | 21993 | - 20043 | glacial | 2.816 | -1.554 | -0.514 | 0.477 | 21993 | 21873 | 4.353 |
| MD02 -2503 | 22147 | - 20197 | glacial | 2.701 | -0.262 | -0.899 | 0.963 | 22147 | 21993 | 2.913 |
| MD02 -2503 | 22288 | - 20338 | glacial | 3.661 | -1.056 | -1.857 | 2.463 | 22288 | 22147 | 3.613 |
| MD02 -2503 | 22414 | - 20464 | glacial | 4.738 | -3.838 | -0.767 | 0.721 | 22414 | 22288 | 5.408 |
| MD02 -2503 | 22499 | - 20549 | glacial | 4.076 | -2.107 | -1.233 | 1.496 | 22499 | 22414 | 3.509 |
| MD02 -2503 | 22611 | - 20661 | glacial | 4.308 | -2.206 | -0.544 | 1.972 | 22611 | 22499 | 2.973 |
| MD02 -2503 | 22724 | - 20774 | glacial | 3.693 | -2.374 | 0.325 | 1.635 | 22724 | 22611 | 4.348 |
| MD02 -2503 | 22850 | - 20900 | glacial | 4.609 | -1.574 | 1.123 | 1.565 | 22850 | 22724 | 3.478 |
| MD02 -2503 | 23005 | - 21055 | glacial | 4.387 | -2.077 | 1.154 | 1.068 | 23005 | 22850 | 1.885 |

| | | | | | | | | | | |
|---|---|---|---|---|---|---|---|---|---|---|
| MD02-2503 | 23145 | -21195 | glacial | 3.389 | -4.298 | 2.491 | -0.109 | 23145 | 23005 | 4.564 |
| MD02-2503 | 23237 | -21287 | glacial | 3.636 | -1.35 | 3.392 | -1.019 | 23237 | 23145 | 4.939 |
| MD02-2503 | 23290 | -21340 | glacial | 0.532 | -1.088 | 1.499 | -1.556 | 23290 | 23237 | 4.376 |
| MD02-2503 | 23472 | -21522 | glacial | 3.491 | -1.017 | 1.341 | 2.203 | 23472 | 23290 | 6.406 |
| MD02-2503 | 23814 | -21864 | glacial | 2.725 | -1.842 | -0.006 | -0.618 | 23814 | 23472 | 6.298 |
| MD02-2503 | 24028 | -22078 | glacial | 2.013 | -2.126 | 0.121 | -1.39 | 24028 | 23814 | 3.83 |
| MD02-2503 | 24135 | -22185 | glacial | 2.944 | -2.367 | -2.09 | -0.923 | 24135 | 24028 | 6.205 |
| MD02-2503 | 24242 | -22292 | glacial | 2.3 | -1.154 | -0.902 | -0.23 | 24242 | 24135 | 3.025 |
| MD02-2503 | 24456 | -22506 | glacial | 1.72 | 0.217 | -1.29 | -0.542 | 24456 | 24242 | 4.02 |
| MD02-2503 | 24617 | -22667 | glacial | 3.388 | -1.47 | -1.903 | -1.073 | 24617 | 24456 | 5.096 |
| MD02-2503 | 25013 | -23063 | glacial | 2.644 | -0.316 | -1.374 | -0.563 | 25013 | 24617 | 2.336 |
| MD02-2503 | 25227 | -23277 | glacial | 2.193 | -0.098 | -2.826 | -1.222 | 25227 | 25013 | 3.312 |
| MD02-2503 | 25441 | -23491 | interglacial | 1.68 | 0.902 | -2.956 | -1.679 | 25441 | 25227 | 3.33 |
| MD02-2503 | 25655 | -23705 | glacial | 2.862 | -1.879 | -3.625 | -1.704 | 25655 | 25441 | 4.836 |
| MD02-2503 | 25869 | -23919 | glacial | 4.254 | -2.673 | -2.655 | -0.244 | 25869 | 25655 | 3.318 |
| MD02-2503 | 26029 | -24079 | glacial | 3.303 | -0.575 | -2.08 | -0.279 | 26029 | 25869 | 3.572 |
| MD02-2503 | 26190 | -24240 | glacial | 1.692 | 0.477 | -2.053 | -0.729 | 26190 | 26029 | 3.845 |

| | | | | | | | | | | |
|---|---|---|---|---|---|---|---|---|---|---|
| MD02-2503 | 26436 | -24486 | glacial | 2.955 | -0.117 | -2.039 | -0.55 | 26436 | 26190 | 2.343 |
| MD02-2503 | 26618 | -24668 | glacial | 2.266 | -0.264 | -1.163 | -0.147 | 26618 | 26436 | 2.511 |
| MD02-2503 | 26832 | -24882 | glacial | 2.503 | 0.255 | -1.94 | -0.116 | 26832 | 26618 | 2.303 |
| MD02-2503 | 27046 | -25096 | glacial | 3.99 | -4.271 | 0.156 | -0.824 | 27046 | 26832 | 6.92 |
| MD02-2503 | 27207 | -25257 | glacial | 3.507 | -0.654 | -1.881 | 0.77 | 27207 | 27046 | 6.446 |
| MD02-2503 | 27378 | -25428 | glacial | 2.505 | -1.813 | 2.801 | 1.226 | 27378 | 27207 | 7.478 |
| MD02-2503 | 27389 | -25439 | glacial | 3.034 | -2.602 | 4.38 | 0.798 | 27389 | 27378 | 3.248 |
| MD02-2503 | 27399 | -25449 | glacial | 2.048 | -2.346 | 4.052 | 1.521 | 27399 | 27389 | 2.689 |
| MD02-2503 | 27410 | -25460 | glacial | 1.653 | -1.493 | 3.551 | -0.318 | 27410 | 27399 | 3.115 |
| MD02-2503 | 27421 | -25471 | glacial | 1.953 | -1.45 | 3.882 | -0.624 | 27421 | 27410 | 2.308 |
| MD02-2503 | 27431 | -25481 | glacial | 1.377 | -1.719 | 3.844 | -1.662 | 27431 | 27421 | 4.604 |
| MD02-2503 | 27442 | -25492 | interglacial | 0.739 | 0.057 | 4.139 | -1.557 | 27442 | 27431 | 4.085 |
| MD02-2503 | 27453 | -25503 | glacial | 1.132 | 0.002 | 2.82 | -2.12 | 27453 | 27442 | 2.474 |
| MD02-2503 | 27463 | -25513 | interglacial | 0.76 | 0.578 | 4.513 | -1.661 | 27463 | 27453 | 2.748 |
| MD02-2503 | 27474 | -25524 | interglacial | 0.219 | 1.397 | 4.479 | -0.299 | 27474 | 27463 | 3.631 |
| MD02-2503 | 27485 | -25535 | glacial | 1.155 | -0.101 | 5.126 | -0.655 | 27485 | 27474 | 3.607 |
| MD02-2503 | 27496 | -25546 | interglacial | 0.514 | 1.627 | 4.113 | -0.959 | 27496 | 27485 | 3.798 |

| | | | | | | | | | | |
|---|---|---|---|---|---|---|---|---|---|---|
| MD02-2503 | 27506 | -25556 | interglacial | -0.082 | 1.441 | 4.352 | -1.493 | 27506 | 27496 | 2.868 |
| MD02-2503 | 27517 | -25567 | interglacial | 0.297 | 2.032 | 4.58 | -1.355 | 27517 | 27506 | 2.364 |
| MD02-2503 | 27528 | -25578 | interglacial | 1.216 | 2.871 | 4.055 | -1.483 | 27528 | 27517 | 2.879 |
| MD02-2503 | 27538 | -25588 | interglacial | -0.206 | 2.253 | 3.912 | -1.955 | 27538 | 27528 | 3.223 |
| MD02-2503 | 27549 | -25599 | interglacial | 0.009 | 1.794 | 6.315 | -0.769 | 27549 | 27538 | 4.521 |
| MD02-2503 | 27560 | -25610 | interglacial | -0.493 | 2.38 | 4.733 | -1.139 | 27560 | 27549 | 2.809 |
| MD02-2503 | 27570 | -25620 | interglacial | 0.012 | 1.937 | 4.774 | -0.788 | 27570 | 27560 | 1.647 |
| MD02-2503 | 27581 | -25631 | interglacial | -1.188 | 1.414 | 5.443 | -1.89 | 27581 | 27570 | 2.967 |
| MD02-2503 | 27592 | -25642 | interglacial | 0.114 | 1.281 | 3.994 | -2.106 | 27592 | 27581 | 2.804 |
| MD02-2503 | 27603 | -25653 | interglacial | 0.231 | 1.489 | 2.889 | -2.119 | 27603 | 27592 | 2.155 |
| MD02-2503 | 27613 | -25663 | interglacial | -0.284 | 1.028 | 4.54 | -1.602 | 27613 | 27603 | 2.945 |
| MD02-2503 | 27624 | -25674 | interglacial | 0.183 | 0.344 | 3.608 | -2.446 | 27624 | 27613 | 2.873 |
| MD02-2503 | 27635 | -25685 | interglacial | 0.317 | 0.645 | 2.378 | -2.169 | 27635 | 27624 | 2.954 |
| MD02-2503 | 27645 | -25695 | glacial | 0.872 | 0.013 | 2.657 | -2.142 | 27645 | 27635 | 2.305 |
| MD02-2503 | 27656 | -25706 | glacial | 1.271 | -1.081 | 2.602 | -1.046 | 27656 | 27645 | 2.721 |
| MD02-2503 | 27667 | -25717 | glacial | 2.662 | -2.504 | 2.55 | -0.806 | 27667 | 27656 | 4.906 |
| MD02-2503 | 27677 | -25727 | glacial | 2.523 | -3.693 | 2.624 | -1.144 | 27677 | 27667 | 2.913 |

| | | | | | | | | | | |
|---|---|---|---|---|---|---|---|---|---|---|
| MD02 -2503 | 27688 | - 25738 | glacial | 3.588 | -3.324 | 2.246 | -1.284 | 27688 | 27677 | 3.093 |
| MD02 -2503 | 27699 | - 25749 | glacial | 3.537 | -3.399 | 1.932 | -1.576 | 27699 | 27688 | 1.663 |
| MD02 -2503 | 27710 | - 25760 | glacial | 2.739 | -2.362 | 1.068 | -0.791 | 27710 | 27699 | 3.86 |
| MD02 -2503 | 27913 | - 25963 | glacial | 3.437 | 0.249 | -1.309 | -0.427 | 27913 | 27710 | 5.449 |
| MD02 -2503 | 28127 | - 26177 | glacial | 4.708 | -0.494 | -2.616 | -0.327 | 28127 | 27913 | 3 |
| MD02 -2503 | 28341 | - 26391 | glacial | 3.21 | 0.246 | -2.28 | -0.05 | 28341 | 28127 | 2.807 |
| MD02 -2503 | 28555 | - 26605 | glacial | 2.664 | 0.421 | -2.781 | -0.38 | 28555 | 28341 | 2.203 |
| MD02 -2503 | 28700 | - 26750 | intergl acial | -2.291 | -0.137 | 3.994 | 0.697 | 28700 | 28555 | 9.784 |
| MD02 -2503 | 28735 | - 26785 | glacial | 0.751 | -1.363 | 3.588 | -0.035 | 28735 | 28700 | 4.806 |
| MD02 -2503 | 28769 | - 26819 | glacial | 2.014 | -0.723 | 0.768 | 1.459 | 28769 | 28735 | 5.809 |
| MD02 -2503 | 28770 | - 26820 | glacial | 2.06 | -5.46 | 3.204 | -0.715 | 28770 | 28769 | 8.986 |
| MD02 -2503 | 28876 | - 26926 | glacial | 1.541 | -1.272 | 1.976 | 2.016 | 28876 | 28770 | 7.894 |
| MD02 -2503 | 28910 | - 26960 | glacial | 4.166 | -2.404 | -3.459 | -3.515 | 28910 | 28876 | 10.059 |
| MD02 -2503 | 28929 | - 26979 | glacial | 2.09 | -0.085 | 3.496 | 2.052 | 28929 | 28910 | 10.057 |
| MD02 -2503 | 28983 | - 27033 | intergl acial | 0.041 | 3.469 | 3.319 | -0.006 | 28983 | 28929 | 6.428 |
| MD02 -2503 | 29026 | - 27076 | intergl acial | -1.359 | 2.542 | 3.869 | -0.666 | 29026 | 28983 | 3.948 |
| MD02 -2503 | 29044 | - 27094 | glacial | 4.192 | -2.767 | -3.041 | -2.375 | 29044 | 29026 | 12.077 |

| | | | | | | | | | | |
|---|---|---|---|---|---|---|---|---|---|---|
| MD02 -2503 | 29121 | - 27171 | glacial | 2.984 | -1.312 | -3.587 | -3.484 | 29121 | 29044 | 3.226 |
| MD02 -2503 | 29422 | - 27472 | glacial | 2.139 | -0.224 | -2.511 | -2.434 | 29422 | 29121 | 3.704 |
| MD02 -2503 | 29562 | - 27612 | glacial | 2.066 | 0.339 | -3.035 | -2.783 | 29562 | 29422 | 1.89 |
| MD02 -2503 | 29689 | - 27739 | glacial | 3.718 | -2.214 | -3.589 | -3.617 | 29689 | 29562 | 4.184 |
| MD02 -2503 | 29864 | - 27914 | intergl acial | 1.901 | 1.343 | -3.792 | -2.316 | 29864 | 29689 | 4.965 |
| MD02 -2503 | 29983 | - 28033 | glacial | 1.662 | 0.536 | -2.269 | -2.252 | 29983 | 29864 | 2.332 |
| MD02 -2503 | 30102 | - 28152 | glacial | 3.737 | -1.241 | -2.452 | -2.464 | 30102 | 29983 | 3.912 |
| MD02 -2503 | 30242 | - 28292 | glacial | 2.217 | -0.409 | -2.284 | -2.087 | 30242 | 30102 | 4.564 |
| MD02 -2503 | 30383 | - 28433 | glacial | 3.378 | -0.282 | -2.338 | -1.33 | 30383 | 30242 | 2.587 |
| MD02 -2503 | 30523 | - 28573 | glacial | 4.626 | -3.723 | 1.174 | 0.314 | 30523 | 30383 | 7.105 |
| MD02 -2503 | 30670 | - 28720 | intergl acial | 2.251 | 1.336 | -2.116 | 0.041 | 30670 | 30523 | 8.516 |
| MD02 -2503 | 30803 | - 28853 | intergl acial | 1.596 | 1.126 | -1.84 | -0.309 | 30803 | 30670 | 2.85 |
| MD02 -2503 | 30894 | - 28944 | intergl acial | 1.448 | 0.611 | -2.452 | -1.165 | 30894 | 30803 | 2.334 |
| MD02 -2503 | 31105 | - 29155 | intergl acial | 1.23 | 1.236 | -2.241 | -0.779 | 31105 | 30894 | 2.367 |
| MD02 -2503 | 31224 | - 29274 | intergl acial | 1.585 | 0.944 | -1.821 | -0.497 | 31224 | 31105 | 1.841 |
| MD02 -2503 | 31364 | - 29414 | intergl acial | 1.277 | 1.289 | -2.15 | -0.454 | 31364 | 31224 | 2.091 |
| MD02 -2503 | 31518 | - 29568 | glacial | 3.348 | 0.5 | -1.277 | 1.596 | 31518 | 31364 | 5.107 |

| | | | | | | | | | | |
|---|---|---|---|---|---|---|---|---|---|---|
| MD02-2503 | 31645 | -29695 | glacial | 3.289 | 0.681 | 3.194 | 0.524 | 31645 | 31518 | 7.501 |
| MD02-2503 | 31715 | -29765 | interglacial | 0.349 | 2.073 | 4.544 | 1.586 | 31715 | 31645 | 5.107 |
| MD02-2503 | 31750 | -29800 | interglacial | 0.813 | 3.355 | 3.324 | -1.574 | 31750 | 31715 | 5.743 |
| MD02-2503 | 31792 | -29842 | interglacial | 0.094 | 2.851 | 4.107 | -1.174 | 31792 | 31750 | 2.68 |
| MD02-2503 | 31820 | -29870 | interglacial | 0.939 | 1.891 | 1.761 | -1.398 | 31820 | 31792 | 3.815 |
| MD02-2503 | 31855 | -29905 | interglacial | -0.276 | 2.488 | 0.42 | -0.173 | 31855 | 31820 | 4.664 |
| MD02-2503 | 32450 | -30500 | interglacial | 0.091 | 2.786 | 2.254 | 0.626 | 32450 | 31855 | 3.235 |
| MD02-2503 | 32520 | -30570 | interglacial | 0.136 | 2.901 | 3.483 | -1.099 | 32520 | 32450 | 4.3 |
| MD02-2503 | 32555 | -30605 | interglacial | -0.38 | 1.18 | 2.278 | -1.342 | 32555 | 32520 | 3.828 |
| MD02-2503 | 32639 | -30689 | interglacial | -0.339 | 1.041 | 0.535 | -3.14 | 32639 | 32555 | 4.875 |
| MD02-2503 | 32667 | -30717 | glacial | 1.041 | -0.053 | 1.265 | -0.415 | 32667 | 32639 | 5.867 |
| MD02-2503 | 32709 | -30759 | glacial | 3.706 | -1.92 | -0.209 | 0.12 | 32709 | 32667 | 6.147 |
| MD02-2503 | 32765 | -30815 | interglacial | 1.55 | 0.795 | -1.303 | -0.838 | 32765 | 32709 | 5.534 |
| MD02-2503 | 32906 | -30956 | interglacial | 2.145 | 1.742 | -2.238 | -0.777 | 32906 | 32765 | 2.292 |
| MD02-2503 | 33046 | -31096 | interglacial | 2.064 | 1.259 | -1.634 | -0.504 | 33046 | 32906 | 2.664 |
| MD02-2503 | 33186 | -31236 | interglacial | 1.2 | 1.564 | -1.648 | -0.611 | 33186 | 33046 | 3.272 |
| MD02-2503 | 33326 | -31376 | glacial | 2.584 | 0.166 | -1.654 | 0.181 | 33326 | 33186 | 3.577 |

| | | | | | | | | | | |
|---|---|---|---|---|---|---|---|---|---|---|
| MD02-2503 | 33467 | -31517 | glacial | 3.47 | -0.574 | 1.658 | 0.711 | 33467 | 33326 | 5.879 |
| MD02-2503 | 33502 | -31552 | glacial | 2.089 | -1.649 | 4.099 | 2.231 | 33502 | 33467 | 5.054 |
| MD02-2503 | 33509 | -31559 | glacial | 1.555 | 0.479 | 4.563 | 1.149 | 33509 | 33502 | 3.84 |
| MD02-2503 | 33516 | -31566 | glacial | 1.158 | 0.074 | 3.886 | 1.113 | 33516 | 33509 | 1.981 |
| MD02-2503 | 33523 | -31573 | glacial | 1.277 | 0.159 | 5.27 | 1.009 | 33523 | 33516 | 3.194 |
| MD02-2503 | 33530 | -31580 | glacial | 1.719 | 0.498 | 3.468 | 1.028 | 33530 | 33523 | 3.051 |
| MD02-2503 | 33537 | -31587 | glacial | 2.208 | -0.888 | 4.485 | 1.108 | 33537 | 33530 | 3.154 |
| MD02-2503 | 33544 | -31594 | interglacial | 1.004 | 0.775 | 3.426 | 0.798 | 33544 | 33537 | 3.408 |
| MD02-2503 | 33551 | -31601 | interglacial | 1.037 | 1.79 | 3.559 | 0.088 | 33551 | 33544 | 2.954 |
| MD02-2503 | 33558 | -31608 | interglacial | 1.007 | 0.791 | 3.221 | -0.106 | 33558 | 33551 | 2.183 |
| MD02-2503 | 33565 | -31615 | interglacial | 1.34 | 0.928 | 1.366 | 0.175 | 33565 | 33558 | 3.351 |
| MD02-2503 | 33572 | -31622 | glacial | 2.029 | 0.541 | 2.754 | 0.734 | 33572 | 33565 | 3.272 |
| MD02-2503 | 33579 | -31629 | interglacial | 2.343 | 2.08 | 0.184 | 0.459 | 33579 | 33572 | 5.055 |
| MD02-2503 | 33586 | -31636 | interglacial | 1.137 | 1.936 | 0.476 | 0.829 | 33586 | 33579 | 3.32 |
| MD02-2503 | 33593 | -31643 | interglacial | 0.614 | 2.149 | 1.132 | 0.948 | 33593 | 33586 | 2.019 |

**Table S3.**

Distance outliers, ages, and reconstructed conditions in the CCS and SBB.

| Calendar Year (AD) | group | Climate event | CCS Conditions | distance | MDS1 | MDS2 |
|---|---|---|---|---|---|---|
| 2006 AD | Anthropocene | | Stratified, warm, bottom-water deoxygenation (1, 2) | 8.955 | -9.617 | -3.089 |
| 1798 AD | Anthropocene | | OMZ relaxation following late LIA cooling and deoxygenation (2) | 15.303 | -1.408 | 2.831 |
| 1605 AD | interglacial | Little Ice Age | Cooling, less oxygenated source water, OMZ intensification (2) | 9.205 | -7.485 | -0.013 |
| 1367 AD | interglacial | Little Ice Age | Cooling, more oxygenated source water, OMZ relaxation; enhanced localized export productivity (2) | 9.041 | -1.045 | 2.16 |
| 10.22 ka | interglacial | pre-Boreal warming | Stratified, warm, increased productivity (3) | 8.576 | -2.108 | 4.055 |
| 14.53 ka | glacial | Bølling warming | Warming of 7˚C in < 500 yrs (4, 5) | 9.517 | 0.902 | -3.929 |
| 14.56 ka | glacial | Bølling warming | | 10.328 | 1.595 | 0.382 |
| 28.7 ka | interglacial | GI-4 | Warm, reduced productivity (3) | 9.784 | -2.291 | -0.137 |
| 28.77 ka | glacial | GI-4 | | 8.986 | 2.06 | -5.46 |
| 28.91 ka | glacial | GI-4 | | 10.059 | 4.166 | -2.404 |

| | | | | | | |
|---|---|---|---|---|---|---|
| 28.93 ka | glacial | GI-4 | | 10.057 | 2.09 | -0.085 |
| 29.04 ka | glacial | GI-4 | | 12.077 | 4.192 | -2.767 |
| references | 1. Goericke et al. 2015 | 2. Deutsch et al. 2011 | 2. Wang and Hendy 2021 | 3. Hendy et al. 2002 | 4. Pak et al. 2012 | 5. White et al. 2013 |

**Table S4.**

Analysis of variance comparing linear models for MDS1, single-predictor regression, and linear mixed model results.

| ANOVA | | | | | | |
|---|---|---|---|---|---|---|
| Model | Res. df | RSS | Δdf | ΔSS | F | p |
| MDS1 ~ Re/Al + Mo/Al + TOC + MAR + $\delta^{15}N$ + SST | 147 | 536.44 | | | | |
| MDS1 ~ $\delta^{15}N$ + TOC + Mo | 150 | 681.95 | −3 | −145.51 | 13.291 | < 0.001*** |
| MDS1 ~ MO + TOC | 151 | 694.81 | −1 | −12.87 | 3.526 | 0.062 . |
| MDS1 ~ TOC | 152 | 695.19 | −1 | −0.38 | 0.104 | 0.748 |
| MDS1 ~ MO | 152 | 1742.42 | 0 | −1047.23 | | |
| MDS1 ~ $\delta^{15}N$ | 152 | 1776.54 | 0 | −34.11 | | |

Note. Models compared sequentially via ANOVA (Type I SS). Res. df = residual degrees of freedom; RSS = residual sum of squares; Δdf = change in degrees of freedom; ΔSS = change in sum of squares. *** p < 0.001; . p < 0.10.

| Single-predictor linear regression | | | | | | |
|---|---|---|---|---|---|---|
| Predictor / Term | β | SE | t | df | p | $R^2$ |
| TOC Intercept | 7.072 | 0.309 | 22.90 | 323 | < 0.001*** | |
| TOC | −3.485 | 0.148 | −23.49 | 323 | < 0.001*** | 0.631 |

| | | | | | | |
|---|---|---|---|---|---|---|
| Mo/Al Intercept | 2.641 | 0.287 | 9.213 | 339 | < 0.001*** | |
| Mo/Al | −4.017 | 0.357 | −11.24 | 339 | < 0.001*** | 0.271 |
| δ¹⁵N Intercept | 7.858 | 1.832 | 4.289 | 339 | < 0.001*** | |
| δ¹⁵N | −1.098 | 0.255 | −4.311 | 339 | < 0.001*** | 0.052 |

| Linear Mixed Model results (MDS1 ~ TOC + δ¹⁵N + Mo/Al + (1 \| cluster group) | | | | | |
|---|---|---|---|---|---|
| Parameter | Estimate | SE | df | t | p |
| Fixed effects | | | | | |
| Intercept | 5.340 | 2.703 | 2.55 | 1.975 | 0.159 |
| TOC | −1.332 | 0.171 | 320.63 | −7.772 | < 0.001*** |
| δ¹⁵N | −0.592 | 0.120 | 319.16 | −4.947 | < 0.001*** |
| Mo/Al | −0.079 | 0.207 | 319.00 | −0.383 | 0.702 |
| Random effects | Variance | SD | | | |
| Group (intercept) | 19.299 | 4.393 | | | |
| Residual | 1.501 | 1.225 | | | |

| | |
|---|---|
| Model fit | |
| $R^2m$ / $R^2c$ | 0.055 / 0.932 |
| N (observations) | 325 |
| Groups | 3 |

Note. SE = standard error; df = Satterthwaite degrees of freedom; $R^2m$ = marginal $R^2$ (fixed effects only); $R^2c$ = conditional $R^2$ (fixed + random effects). *** $p < 0.001$.

**Table S5.**

Analysis of variance comparing linear models for ecological distance (euclidean dissimilarity) and single-predictor regression outputs.

ANOVA

| Model | Res. df | RSS | Δdf | ΔSS | F | p |
|---|---|---|---|---|---|---|
| distance ~ re + MO + TOC + MAR + $\delta^{15}N$ + temp | 146 | 574.69 | | | | |
| distance ~ $\delta^{15}N$ + TOC + MO | 149 | 589.61 | −3 | −14.921 | 1.264 | 0.289 |
| distance ~ MO + TOC | 150 | 670.42 | −1 | −80.811 | 20.530 | < 0.001*** |
| distance ~ TOC | 151 | 709.35 | −1 | −38.933 | 9.891 | 0.002** |
| distance ~ MO | 151 | 670.82 | 0 | 38.529 | | |
| distance ~ $\delta^{15}N$ | 151 | 678.77 | 0 | −7.951 | | |

Note. Models compared sequentially via ANOVA (Type I SS). Res. df = residual degrees of freedom; RSS = residual sum of squares; Δdf = change in degrees of freedom; ΔSS = change in sum of squares. * $p < 0.01$; *** $p < 0.001$.

Single-predictor linear regression

| Predictor / Term | β | SE | t | df | p | $R^2$ |
|---|---|---|---|---|---|---|
| TOC Intercept | 3.476 | 0.315 | 11.049 | 322 | < 0.001*** | |
| TOC | 0.459 | 0.152 | 3.017 | 322 | 0.003** | 0.027 |

| | | | | | | |
|---|---|---|---|---|---|---|
| Mo/Al Intercept | 3.741 | 0.189 | 19.826 | 338 | < 0.001*** | |
| Mo/Al | 1.072 | 0.238 | 4.505 | 338 | < 0.001*** | 0.057 |
| $\delta^{15}N$ Intercept | −0.216 | 1.049 | −0.205 | 338 | 0.837 | |
| $\delta^{15}N$ | 0.650 | 0.146 | 4.462 | 338 | < 0.001*** | 0.056 |

| Linear Mixed Model results (distance ~ TOC + $\delta^{15}N$ + Mo/Al + (1 \| cluster group) | | | | | |
|---|---|---|---|---|---|
| Parameter | Estimate | SE | df | t | p |
| Fixed effects | | | | | |
| Intercept | −2.101 | 1.407 | 82.39 | −1.493 | 0.139 |
| TOC | 0.334 | 0.238 | 13.12 | 1.403 | 0.184 |
| $\delta^{15}N$ | 0.760 | 0.184 | 202.19 | 4.136 | < 0.001*** |
| Mo/Al | 0.623 | 0.323 | 319.72 | 1.930 | 0.055 . |
| Random effects | Variance | SD | | | |
| Group (intercept) | 0.229 | 0.479 | | | |
| Residual | 3.612 | 1.901 | | | |

| | |
|---|---|
| Model fit | |
| $R^2m$ / $R^2c$ | 0.141 / 0.192 |
| N (observations) | 324 |
| Groups | 3 |

Note. SE = standard error; df = Satterthwaite degrees of freedom; $R^2m$ = marginal $R^2$ (fixed effects only); $R^2c$ = conditional $R^2$ (fixed + random effects). *** $p < 0.001$.

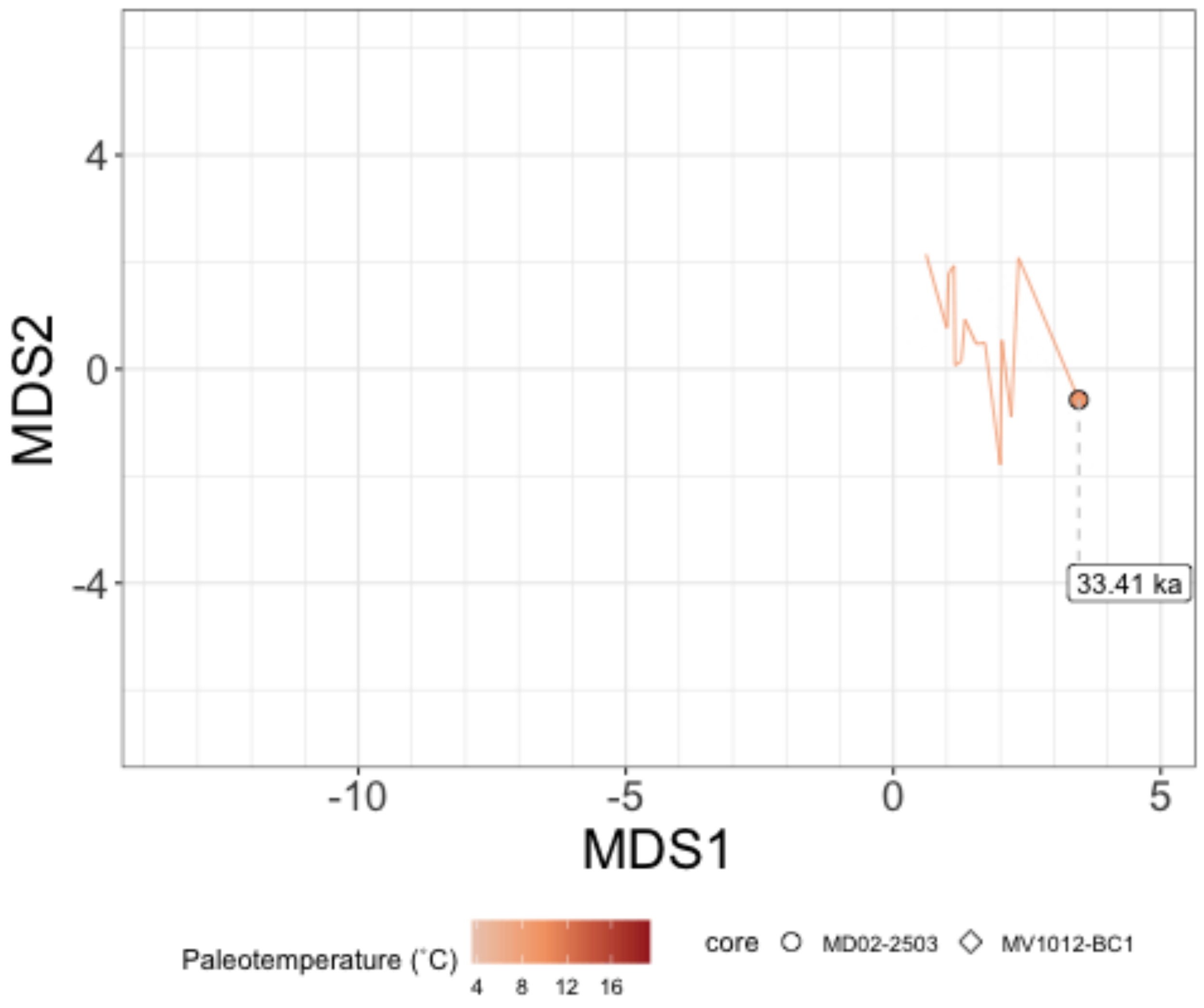


**Movie S1.**
2D NMDS plot showing groupings of rapid warming events. Grey line shows connections between points based on sample age. Color represents SST (same scale as Main Text Figure 1).
